# Evaluating anonymized genome re-identification using polygenic predictions and its implications for data privacy

**DOI:** 10.64898/2026.06.10.731306

**Authors:** Théo Cavinato, Robin J. Hofmeister, Zoltán Kutalik

**Affiliations:** Department of Computational Biology, University of Lausanne, Lausanne, Switzerland; Swiss Institute of Bioinformatic (SIB), University of Lausanne, Lausanne, Switzerland; University Center for Primary Care and Public Health, Lausanne, Switzerland; Estonian Genome Centre, Institute of Genomics, University of Tartu, Tartu, Estonia

## Abstract

Re-identification by phenotypic prediction aims to determine whether a genome belongs to a specific individual by comparing the individual’s known traits with those predicted from the genome. This type of tracing attack is widely discussed in the genomic privacy literature, yet previous studies have been criticized for overstating its practical risks. Over the past decade, genome-wide association studies (GWAS) with increasing sample size improved the accuracy of phenotypic prediction, potentially enhancing such attacks.

To quantify their real-world threat, we developed a probabilistic framework that estimates the likelihood of a match between an individual’s observed traits and polygenic scores (PGS) derived from a genome, while accounting for prediction accuracy and genetic and environmental correlations between the traits. We benchmarked this re-identification method and examined how the *prior* probability (reflecting the *a priori* chance that a random genome and set of traits correspond to the same person) affects performance. Finally, we assessed whether sensitive information could be inferred through this attack by attempting to predict multiple sensitive haplotypes, such as APOE-*ε*4 (linked with Alzheimer’s disease).

Our re-identification method outperformed a state-of-the-art tool, and reached a precision above 99% for a recall of 40% when considering a *prior* of 50%. However, after considering real-world settings, we estimated that realistic *priors* would not exceed 4 *×* 10^−4^%, resulting in a precision lower than 0.13% at the same recall (40%). The inference of sensitive genotypes also proved ineffective, as achieving a precision above 50% for identifying APOE-*ε*4 carriers was only possible at a recall below 20%. To conclude, although re-identification by phenotypic prediction is technically feasible, our findings indicate that its effectiveness in real-world conditions is limited. These results counterpoint to earlier claims of severe genomic privacy risks and offer guidance for policymakers, biobank administrators, and research participants.

## Introduction

Human genome sequences are often considered sensitive data as they can reveal personal information about their owners; with access to a genome and publicly available Genome-wide association study (GWAS) summary statistics, it is possible to infer an individual’s ancestry, his relatedness to others and their risk of developing certain diseases. The misuse of such inferred information may result in genetic discrimination. As a consequence, many countries have regulated the use of genetic data, e.g. with the Genetic Information Nondiscrimination Act (GINA) in the United States, the Genetic Non-Discrimination Act in Canada, GDPR and national laws in European countries, and the Bioethics and Safety act in South Korea [1, 2].In other countries, however, regulations on the use of genetic data remain limited. In New Zealand, for example, insurance companies can request and use the results of previous genetic tests, even for health insurance underwriting [3].Most countries regulating the use of genetic data prohibit its use in health insurance. However, these regulations are often more flexible when it comes to life insurances. This is the case in Germany, Israel, the Netherlands, Sweden, Switzerland, and the United Kingdom, where insurers can use genetic information for underwriting policies that exceed a certain threshold. For example, in the United Kingdom, insurance companies may request the results of a predictive genetic test for Huntington’s disease when the policy value exceeds £500 000 [4].But genetic discrimination can go beyond insurance companies. In 2021, the *European Society of Human Genetics* wrote an article to warn against the misuse of genetic data for discrimination purposes [5].The authors described how different countries attempted or succeeded in imposing compulsory DNA sequencing, and how this practice might lead to discrimination against minorities.

The fear of being the target of genetic discrimination affects scientific research, as it discourages people from participating in genetic studies [1, 4].To protect participants’ privacy, genetic data in research cohorts are usually de-identified (i.e. personal identifiers are removed). However, in 2014, Erlich and Narayanan outlined several approaches by which the owner of a de-identified genome could potentially be re-identified [6].They referred to these approaches as *identity tracing attacks*, which covers several distinct strategies: (1) **Tracing with metadata**, which involves using contextual data (e.g. e-mail address or zip code) published alongside the genome to re-identify the individual; (2) **Tracing by genealogical triangulation**, which identifies genetic relatives in order to infer the identity of the genome’s owner; (3) **Tracing via side-channel leaks**, which exploits unintended data leakage that could render a genome identifiable; and (4) **Tracing by phenotypic prediction**, which compares phenotypes predicted from the genome to known phenotypes of individuals in order to establish a match. At the time of writing, the first three types of attack had either been demonstrated in real-world scenarios or were considered feasible in certain situations. In contrast, tracing by phenotypic prediction was deemed unlikely, as the genetic knowledge available then explained only a small proportion of the variability in a limited number of phenotypes. Nonetheless, a year later, Humbert et al. proposed an approach to de-anonymize genetic datasets based on this idea [7].They considered a scenario where they knew that an individual was part of a genetic dataset, and their goal was to determine which genome corresponded to that individual. However, their method faced the same limitation highlighted by Erlich and Narayanan: only a small number of phenotypes (10) could be used, and their poor predictability from DNA variants inevitably led to weak performance. Moreover, the evaluation of their method was based on re-identifying an individual among 80 other individuals, which is not practically realistic with available datasets that are much larger and typically contain tens of thousands of individuals. Later works on re-identification by phenotypic prediction tried to include face morphology into the prediction, such as the work of Lippert et al. [8] who used it alongside age, sex, height, and voice signature. However, their paper was heavily criticized by Yaniv Erlich, who, among other critics, emphasized that facial morphology contributed poorly to the re-identification process, which was primarily driven by sex and ancestry principal component coordinates of the genomes [9].In the wake of this work, research on re-identification of genetic data has shifted to the use of images to re-identify individuals [10].

Despite some flaws in previous studies, re-identification by phenotypic prediction is still cited as a major threat in reviews covering genetic privacy [11, 12].During the last decade, Genome-wide association studies (GWAS) deepened our understanding of the genetic bases of complex traits and gradually increased the predictive performance of polygenic predictors for thousands of phenotypes. The summary statistics of these studies can be freely accessed online, allowing anyone with a genome to predict its phenotypes through PolyGenic Scores (PGSs). Here, using summary statistics computed with state-of-the-art methods and simulations based on data from the UK Biobank, one of the largest population cohorts worldwide, we evaluated the performance of re-identification by phenotypic prediction in different real-world settings, in order to assess the actual risk posed by this approach. To do so, we developed a method that compares the observed phenotypes of an individual to the predicted phenotypes from a genome, and used this match score to predict whether the genome belongs to the same individual (match) or not (mismatch). Finally, we assessed how this method could be applied in different contexts: (i) verifying an individual’s claim to be the owner of a genome based on their phenotypes, (ii) determining whether an individual is part of a biobank of genomes using their phenotypes, and (iii) inferring an individual’s sensitive genotypes from their phenotypes. Complying with UK Biobank’s Material Transfer Agreement and UK data protection law, all re-identification attempts are simulated, and no participant data were exposed or at risk in the process.

## Results

### Overview of the method

#### Box 1

Re-identification using polygenic predictions

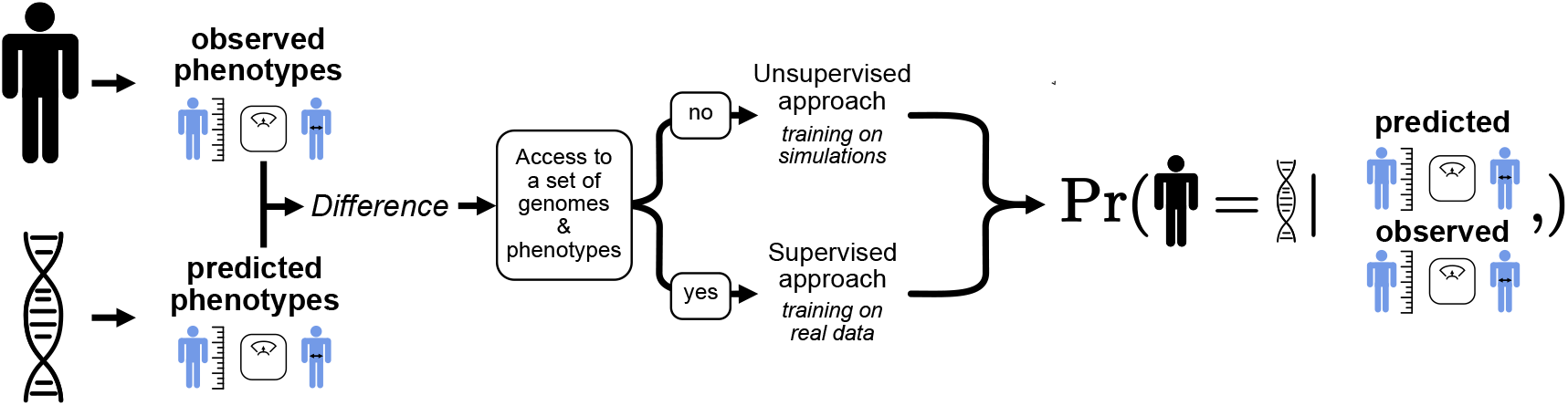

An adversary seeks to determine whether a genome of interest belongs to a known individual. To do so, the adversary uses the individual’s observed phenotypes and infers the corresponding phenotypes from the genome. It then computes the *difference* between the **observed** and **predicted** phenotypes. This difference is evaluated against its expected distribution under two competing hypotheses: that the genome belongs to the individual and that it does not. These distributions can be derived solely from summary statistics (**unsupervised approach**) or, when a training dataset of paired genomes and phenotypes is available, estimated from such data (**supervised approach**). Through this final evaluation, the adversary obtains a quantitative probability that the genome matches the individual.

At a high level, our method quantifies the concordance between the observed phenotypes of an individual and the phenotypes predicted from a genome in order to infer the probability of a match (**Box 1**). More formally, based on *T* traits of an individual *I* and the corresponding *T* PGSs of a genome *G*, our method aims to distinguish between two scenarios: genome *G* belongs to individual *I* (*H*_1_) and *G* does not belong to *I* (*H*_0_) (**Figure 1**). To compare *G* to *I*, we use a log-likelihood ratio (*LLR*) defined as the ratio of the probability of observing *y* (the vector of *I*’s phenotypes) given *s* (the vector of *G*’s PGSs) and *H*_1_ to the probability of observing *y* given *H*_0_, taking into account the pairwise environmental correlations between the traits (Σ_*e*_) and the proportion of the trait variance explained by the PGSs (*r*^2^) (**Figure 1A - LLR calculation**).

**Figure 1:**
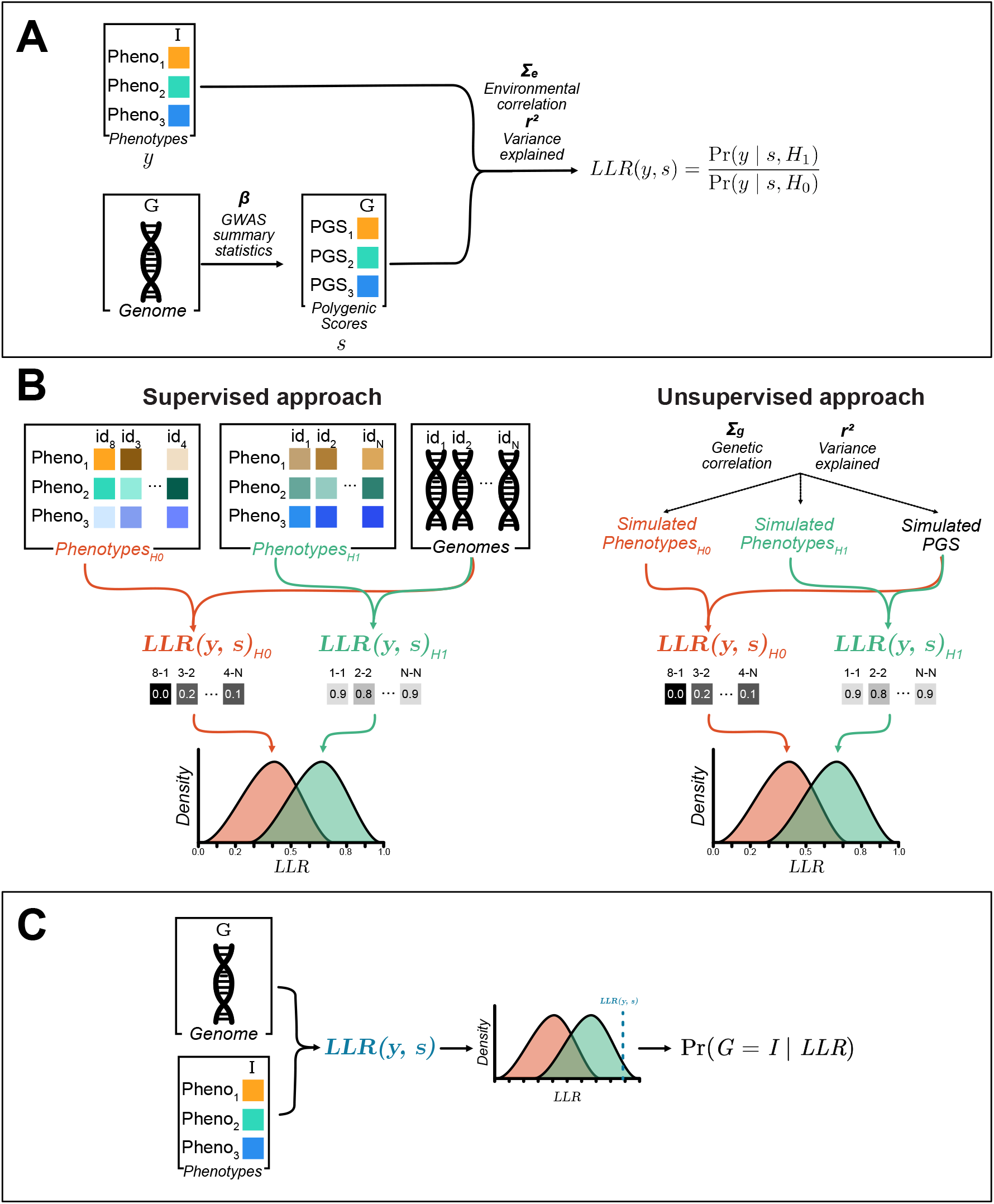
Method description. Our method estimates the probability that a genome *G* belongs to an individual *I*, based on the know phenotypes of *I* and the corresponding polygenic scores (PGSs) of *G*. **A. LLR calculation**. The comparison between PGSs (*s*) and phenotypes (*y*) is performed using a log-likelihood ratio, *LLR*(*y, s*), which accounts for the variance explained by the PGS in each traits (*r*^2^) and the environmental correlation between them (Σ_*e*_). **B. Model training**. To learn the distribution of the *LLR* under the hypotheses that *G* belongs to *I* (*H*_1_) or does not belongs to *I* (*H*_0_), our method simulates pairs of matching and mismatching genomes and phenotypes and compute their *LLR* values. This can be done using a supervised approach if a dataset of genotypes and phenotypes is available, or using only summary statistics (unsupervised approach) if such individual-level data is not accessible. **C. Model application**. The observed *LLR* between *G* and *I* is then compared to the learned distributions to estimate probability that *G* belongs to *I*.

An adversary aiming to guess whether a genome of interest belongs to an individual of interest would first need to learn the distribution of this *LLR* under *H*_0_ and *H*_1_. These distributions can be learned either from summary statistics (**unsupervised approach**) or using, if available, a representative dataset of genomes and individuals (**supervised approach**) (**Figure 1B - Model training**). The **supervised approach** represents an upper limit of re-identification performance, while the **unsupervised approach** is more realistic as it only requires summary statistics that are publicly available.

Once these distributions are learned, the adversary can compute the *LLR* between a genome and an individual of interest, compare it to the learned *LLR* distributions under *H*_0_ and *H*_1_, and derive the probability that genome *G* belongs to individual *I* (Pr(*G* = *I* |*LLR*) (**Figure 1C - Model application**). We describe the method in full details in the **Materials and Methods** section and its software implementation, termed **PGMatch** (**P**henotype-**G**enotype **M**atch using PGS), is freely available on GitHub.

### Effect of the number of phenotypes

To assess the effect of the number of traits on the re-identification performance of our method, we first investigated the distribution of the *LLRs* expected by our model (either based on summary statistics or on a representative dataset of genomes and individuals (**Figure 1 B**)) to the distribution observed in the population (**Figure 1C**). We selected 76 phenotypes from the UK biobank that were present in *>* 100 000 individuals of each sex. To avoid redundancy in the information they bring, when two phenotypes had a correlation *>* 0.8, we only kept the phenotype whose variance explained by the PGS was higher (for the list of the resulting phenotypes and the corresponding variance explained, see **Table S1**). This resulted in 59 traits, out of which we selected the 40 for which the PGS explained the most trait variance (0.35 *≥r*^2^ *≥*0.05). We applied both the **supervised approach** (**Figure 2 A**) and **unsupervised approach** (**Figure 2B**) using these 40 phenotypes, and 1 000 matching PGSs-traits pairs and 1 000 mismatching pairs as a test set. For the supervised approach, we assume that we had 1 000 individuals with genomes and phenotypes available that were used to train the model. These individuals were independent from and non-overlapping with those used in the test set.

**Figure 2:**
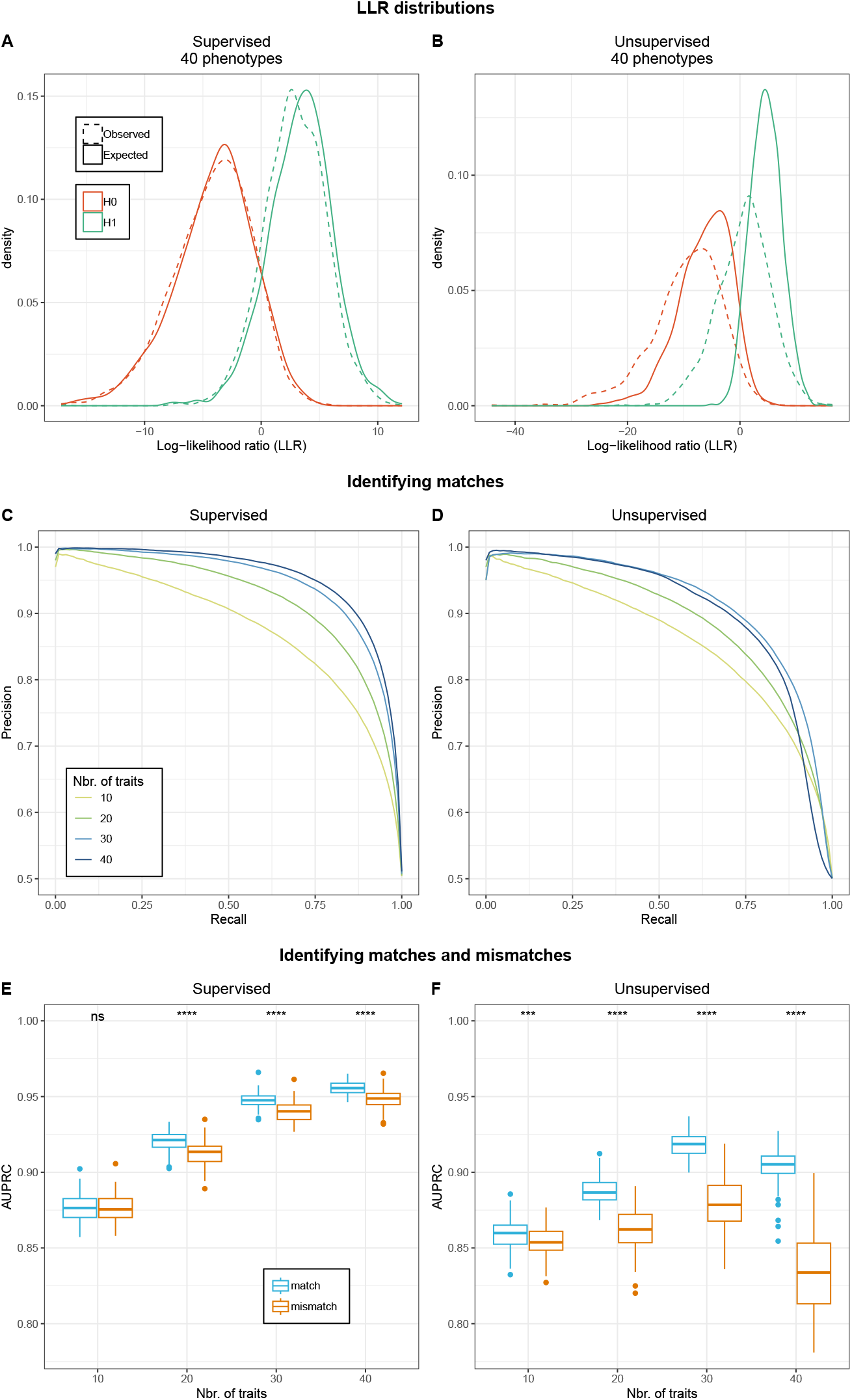
Effect of the number of phenotypes on our approach. **A.,B**. Distribution of the *LLRs* values expected from summary statistics or a train set (solid lines) and observed in a test set (dashed lines) when comparing a genome to its actual traits (*H*_1_ in green) and to the traits of a randomly sampled other individual (*H*_0_ in red). **A**. and **B**. show the results estimated by the supervised and unsupervised approach, respectively. **C**., **D**., Precision and recall of our inference of matches averaged across 100 experiments. Colors indicate the number of traits used for inference. **E**., **F**. Comparison of the area under the precision–recall curve (AUPRC) for match (blue) and mismatch (orange) inference using different number of phenotypes.

Using the unsupervised approach, the expected distributions matched poorly the observed distributions, except for high *LLR* values. With the supervised approach, we observe a strong match in the *LLR* distribution in the training- and the test set. This strong agreement was unsurprising as the expected distribution of the *LLR* in the supervised approach is derived from the training set and the samples in the test and training sets are randomly drawn from the same population. Moreover, the observed overlap between the likelihood values under *H*_0_ and *H*_1_ was much stronger in the unsupervised- than in the supervised approach. We observed the same patterns when repeating the experiment using 10, 20, and 30 phenotypes (**Supp. Fig. S1**). Finally, we observed a stronger distinction between the *H*_0_ and *H*_1_ distributions as the number of phenotypes increased.

Next, we investigated the effect of the number of phenotypes on the inference of match- and mismatch probabilities (*Pr*(*G* = *I* |*LLR*) *vs Pr*(*G* ≠ *I* |*LLR*)). To this end, we transformed the *LLR* values computed in the first step into *Pr*(*G* = *I* |*LLR*) and *Pr*(*G* ≠ *I* |*LLR*) (as described in **Figure 1C**) and assessed the precision and recall we would obtain by using different thresholds for the classification. We repeated the experiment 100 times, each time randomly sampling new training and testing sets. In (**Figure 2C,D**) we report precision–recall curves for the inference of matches, constructed by averaging precision values at each recall level across all experiments for the same number of phenotype. As expected, based on the distribution of the *LLR* values, the supervised approach performed better than the unsupervised as shown by its higher area under the precision–recall curve (AUPRC) at equal number phenotypes (**Table 1**). In both scenarios, the number of phenotypes increased the performance. In (**Figure 2E,F**) we compared the inference of match to the inference of mismatch across the 100 resampled data sets. In both scenarios, for experiments performed with 20 or more phenotypes, match-inference performed better than mismatch-inference, as shown by the significantly higher AUPRC (one-sided t-test) for inferring matches.

**Table 1.** Average area under the precision-recall curve (AUPRC) across 100 experiments for matches and mismatches inference depending on the approach and number of phenotypes.

| Number of traits | Inference type | Avg. AUPRC supervised | Avg. AUPRC unsupervised |
| --- | --- | --- | --- |
| 10 | match | 0.8761 | 0.8589 |
| 20 | match | 0.9203 | 0.8873 |
| 30 | match | 0.9464 | 0.9159 |
| 40 | match | 0.9558 | 0.9044 |
| 10 | mismatch | 0.8761 | 0.854 |
| 20 | mismatch | 0.9123 | 0.8619 |
| 30 | mismatch | 0.9385 | 0.8804 |
| 40 | mismatch | 0.9492 | 0.8280 |

Interestingly, the prediction accuracy of the unsupervised approach decreased when increasing the number of phenotypes to 40. To investigate this effect, we examined the AUPRC when incrementally adding phenotypes from 30 to 40 (**Supp. Fig. S2**). We identified two instances where performance dropped rather than improved with the unsupervised approach, corresponding to the addition of the 31st (BMI) and 32nd (immature reticulocyte count) phenotypes. To further explore this, we excluded these two phenotypes and repeated the analysis from 30 to 38 phenotypes (**Supp. Fig. S3**). In this setting, no performance decrease was observed. We therefore hypothesize that these drops arise from non-linear interactions or residual dependencies between phenotypes that are not adequately captured by the unsupervised model. For instance, BMI is related to traits such as standing height or whole-body water mass in a non-linear manner. Similarly, immature reticulocyte count is biologically linked to other reticulocyte-related traits already included in the model, such as mean reticulocyte volume and high light scatter reticulocyte count. These observations highlight that accurately modeling the correlation structure between phenotypes is challenging, and that poor estimation can reduce the performance of the unsupervised approach.

Finally, we further investigated what might explain the underperformance of the unsupervised compared to the supervised approach. As mentioned above, the unsupervised approach relies only on summary statistics, whereas the supervised approach estimates the environmental correlation and PGS correlation from a training set of genomes and phenotypes. In contrast, the unsupervised approach assumes these quantities to be equal to the genetic correlation estimated from summary statistics. To investigate the impact of this approximation, we plugged in the environmental correlations estimated by the supervised model into the unsupervised model (Unsupervised + CE, **Supp. Fig. S4**). The performance of the unsupervised approach improved when this information was included, highlighting the substantial differences between the genetic correlation matrix estimated from summary statistics and the environmental correlation matrix derived from observed data. However, even with the knowledge of the environmental correlation, the unsupervised approach still did not reach the performance of the supervised approach. This result is expected, as the supervised method learns the environmental and PGSs correlations, as well as the variance explained by each PGS, directly from individual-level data, whereas the unsupervised approach can only approximate these quantities from summary statistics.

### Impact of the size of the training set

To investigate the effect of the size of the training set on the **supervised approach**, we repeated the experiment with 40 phenotypes but using 100, 200, 300, 400, 500, 1 000 or 5 000 individuals in the training set. As expected, AUPRC increased with the number of individuals (**Figure 3**, dark blue), but quickly plateaued at 500 training samples, where we already reached an average AUPRC of 0.9516 for matches and 0.9426 for mismatches, which is very close to what would be obtained with 5 000 individuals (AUPRC match=0.9586, AUPRC mismatch=0.9519). We repeated the experiment using 10, 20 and 30 phenotypes and observed the same patterns (**Figure 3**, yellow, green, light blue). As expected, the variability of the AUPRC across samplings decreases as the number of individuals increases (using 40 phenotype, for match-inference sd = 0.0123 at 100 individuals vs sd = 0.0034 at 5 000 and for mismatch-inference sd = 0.0335 at 100 individuals vs sd = 0.0053 at 5 000).

**Figure 3:**
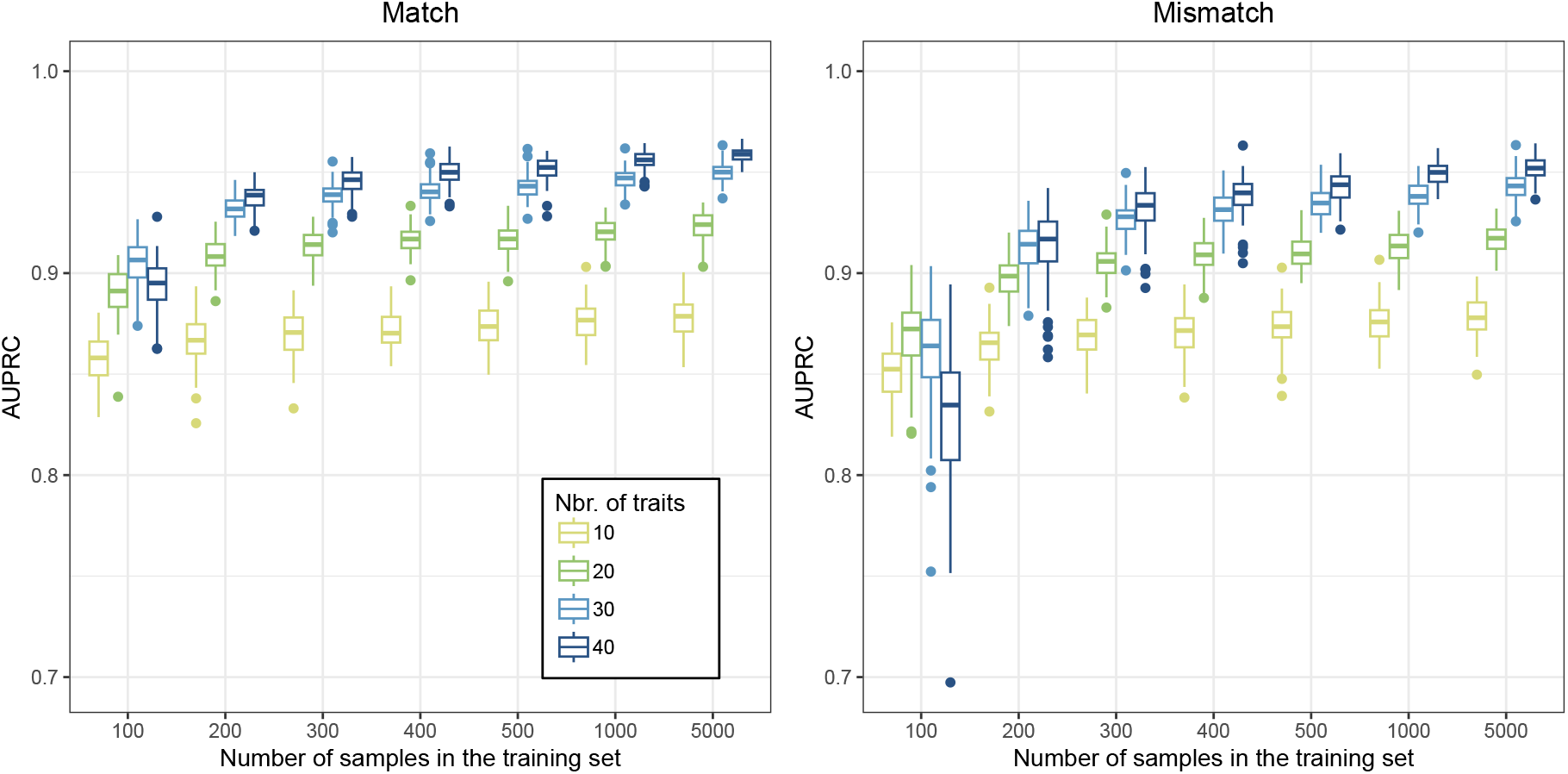
Effect of the size of the training set on our approach. AUPRC (area under the precision-recall curve) depending on the number of individuals in the training set (x-axis) across 100 experiments. Colors correspond to the number of phenotypes used in the experiment. Left: AUPRC for the inference of matches. Right: AUPRC for the inference of mismatches.

### Choices of prior match probabilities

The previous sections demonstrated that the method performs well when assessing its performance on an equal number of matches and mismatches. However, in a real world setting, when one randomly samples a genome and an individual from a population, the *a priori* probability of a match is much lower than that of a mismatch. For instance, when sampling any UK resident and any genome from the UK Biobank, the probability that they are a match depends on the probability for the individual to be part of the biobank and the probability for the genome to be his i.e. 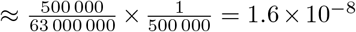. We call it prior (A).

One could make use of the recruitment criteria to increase this probability: UK Biobank participants were selected from individuals living within a 40km radius of a UK Biobank assessment centre and who were between 40 and 69 years of age between 2006 and 2010. Using national statistics and openly accessible data of the UK Biobank, we can define the probability of an individual to be part of the UK Biobank sample depending on the area they leaved in (see **Materials and Methods** for more details). For instance, when randomly sampling a male individual from Bristol and comparing him with a genome from the UK Biobank, the probability that they belong to the same person (a match) is 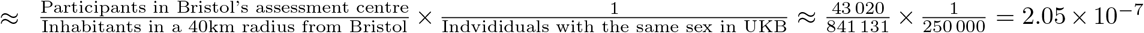 (see **Supp. Tab. S2** for analogous probabilities for each assessment centre in England and Wales). We call it prior (B).

Another way to increase this probability is to assess whether the individual belongs to the biobank. Solely using phenotype-genome matching, one cannot tell if a sample with a given phenotype is part of a biobank (see **Supplementary Note 1**). However, the adversary might have learnt it through another method (e.g. social engineering). In such a case, the fraction of matches would increase to 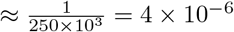. We call it prior (C).

### Precision-recall curves adapted to realistic priors of matching probability

To assess the effect of the prior on the precision of our method, we repeated the re-identification experiment using 40 phenotypes, this time with unbalanced sets of matches and mismatches as test sets. To this end, we generated datasets containing 100 matches combined with 100 to 10^8^ mismatches, as well as 100 mismatches combined with 10^3^ or 10^4^ matches, and estimated the corresponding precision and recall for each dataset. These represent match-to-mismatch ratios ranging from 10^−5^ to 10^2^. We repeated this procedure 100 times, resampling new datasets at each iteration, and report in **Figure 4A-B** the average precision at different recall levels obtained for each match:mismatch ratio. When we originally assumed a *prior* probability *>* 50% for a phenotype profile and genome to be a match, the AUPRC values were well above 0.9 with both methods. When we decreased this *prior* to 10^−1^, the AUPRC dropped to 0.7822 and 0.6378 with the supervised and unsupervised approaches, respectively. (**Figure 4A-B** and **Table 2**). Using a prior of the same order of magnitude as prior (C) (dark blue color **Figure 4A-B**), the AUPRC further dropped to 0.0030 and 0.0009 for the supervised and unsupervised method, respectively. The effect of priors (A) and (B) would result in an even lower AUPRC than prior (C), the computation of which would have been pointless waste of CPU power.

**Table 2.**
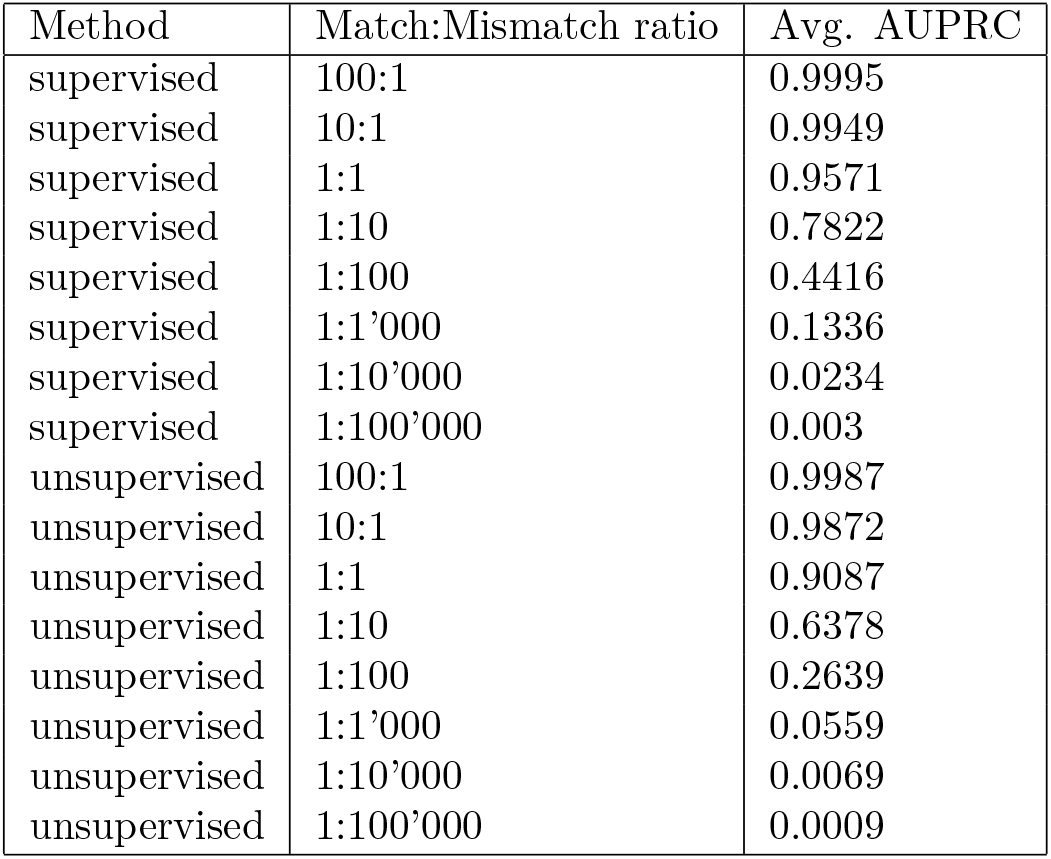
Average area under the precision–recall curve (AUPRC) for matches inference across 100 experiments depending on the method and Match:Mismatch ratio.

| Method | Match:Mismatch ratio | Avg. AUPRC |
| --- | --- | --- |
| supervised | 100:1 | 0.9995 |
| supervised | 10:1 | 0.9949 |
| supervised | 1:1 | 0.9571 |
| supervised | 1:10 | 0.7822 |
| supervised | 1:100 | 0.4416 |
| supervised | 1:1'000 | 0.1336 |
| supervised | 1:10'000 | 0.0234 |
| supervised | 1:100'000 | 0.003 |
| unsupervised | 100:1 | 0.9987 |
| unsupervised | 10:1 | 0.9872 |
| unsupervised | 1:1 | 0.9087 |
| unsupervised | 1:10 | 0.6378 |
| unsupervised | 1:100 | 0.2639 |
| unsupervised | 1:1'000 | 0.0559 |
| unsupervised | 1:10'000 | 0.0069 |
| unsupervised | 1:100'000 | 0.0009 |

**Figure 4:**
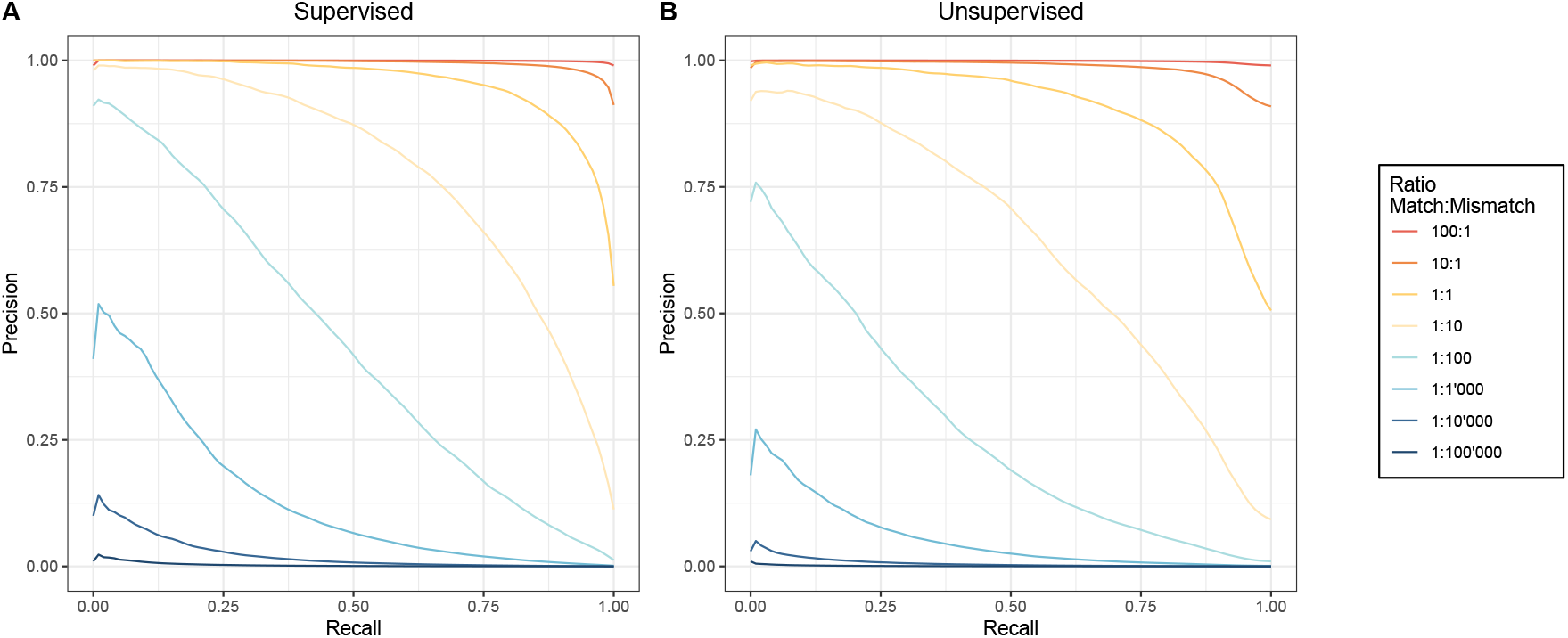
Re-identification precision–recall with uneven number of matches and mismatches. Effect of the Match:Mismatch ratio on the precision-recall curves of re-identificaiton by phenotypic prediction. Warmer colors indicate more matches than mismatches; cooler colors indicate the opposite. **A** and **B** show results for the supervised and unsupervised approaches, respectively.

### Evaluation against the closest existing method

IDEFIX [13] is a method designed to identify sample mix-ups in biobanks by comparing individuals’ phenotypes to polygenic scores (PGSs) derived from their genomes, assuming a known mix-up rate. Briefly, for each phenotype, it fits a linear regression between PGSs and phenotypes and estimates the distribution of residuals when comparing a genome to its true phenotypes versus those of a randomly sampled individual. However, IDEFIX does not directly provide a mix-up–rate–agnostic matching probability between a genome and a phenotype profile. In principle, the LLRs it estimates could be used as input to our method. Nonetheless, the resulting performance is expected to differ, as the two approaches diverge along three key aspects.

The first key difference between the two methods lies in the algorithm used to estimate the *LLR*. IDEFIX computes *LLRs* for all pairs of individuals in the dataset, resulting in *𝒪*(*n*^2^) time and memory complexity. In contrast, PGMatch computes LLRs only for the input phenotype–PGS pairs, yielding 𝒪 (*n*) scaling. We evaluated computational time and memory usage as a function of the number of phenotypes and individuals. We first simulated datasets with 10–40 phenotypes and 1 000 training and testing individuals, and applied both methods across 10 replicates. IDEFIX’s runtime and memory increased with the number of phenotypes and remained consistently higher than PGMatch (**Supp. Fig. S5 A,B**). We then fixed the number of phenotypes to 10 and varied the number of individuals in the testing set from 1 000 to 10 000. IDEFIX exhibited quadratic scaling with dataset size (**Supp. Fig. S5 C,D**). and, across all settings, PGMatch consistently outperformed IDEFIX in both time and memory usage with differences reaching up to two orders of magnitude.

The second key difference is that IDEFIX does not weight phenotypes by the proportion of their variance explained by the PGS (*r*^2^). To assess the impact of this, we simulated phenotypes and PGSs under varying *r*^2^ configurations: (i) *r*^2^ = 0.5 for all phenotypes; (ii) *r*^2^ = 0.05 for all phenotypes; (iii) a mixed setting with half of the phenotypes at *r*^2^ = 0.5 and half at *r*^2^ = 0.05; and (iv) a highly heterogeneous setting with one phenotype at *r*^2^ = 0.5 and the remaining phenotypes at *r*^2^ = 0.05. In all scenarios, we considered 10 phenotypes and datasets of 1 000 individuals for training and 1 000 for testing, with no environmental or PGS correlations between traits. We applied both PGMatch and IDEFIX to the simulated data. Each scenario was repeated 100 times, and results were averaged (**Supp. Fig. S6 C,D**). Across all settings, PGMatch consistently outperformed IDEFIX. The largest performance gap occurred under heterogeneous architectures, where one phenotype had high heritability (*r*^2^ = 0.5) and the others had low heritability (*r*^2^ = 0.05).

The third key difference is that IDEFIX does not account for environmental correlations between phenotypes. We evaluated the effect of correlation between the PGS (Σ_*s*_) and environmental (Σ_*e*_) correlations through simulations. We considered five scenarios: (i) Σ_*e*_ = 0, Σ_*s*_ = 0; (ii) Σ_*e*_ = 0, Σ_*s*_ = 0.5; (iii) Σ_*e*_ = 0.5, Σ_*s*_ = 0; (iv) Σ_*e*_ = 0.5, Σ_*s*_ = 0.5; and (v) a heterogeneous setting where Σ_*e*_ = Σ_*s*_ = 0.5 for half of the phenotypes and Σ_*e*_ = Σ_*s*_ = 0 for the other half. All simulations used 10 phenotypes and a training set of 1 000 individuals and a testing set of 1 000 individuals. *r*^2^ values mimicked those of the 10 best-predicted phenotypes in our real data application (UK Biobank). We applied both PGMatch and IDEFIX to each simulated dataset. PGS correlations had little impact on the relative performance of the two methods (**Supp. Fig. S7**). In contrast, environmental correlations substantially deteriorated the performance of IDEFIX relative to PGMatch.

Another crucial difference is that our PGMatch approach has an unsupervised version that works using summary statistics only, while IDEFIX always requires raw training data for matched genotypes and phenotypes. This makes our method much more amenable to realistic adversary situations.

Finally, we repeated the experiment under realistic priors using IDEFIX and compared the results to PGMatch. Specifically, we considered match-to mismatch ratios from 10^−2^ to 10^2^, using 40 phenotypes. Each setting was repeated 100 times and results were averaged. Due to computational constraints, IDEFIX could not be evaluated beyond 10^4^ matches or mismatches. PGMatch consistently outperformed IDEFIX across all settings **Figure 5**. The performance gap was modest when matches were relatively abundant, but increased sharply as mismatches dominated, becoming substantial under highly imbalanced regimes.

**Figure 5:**
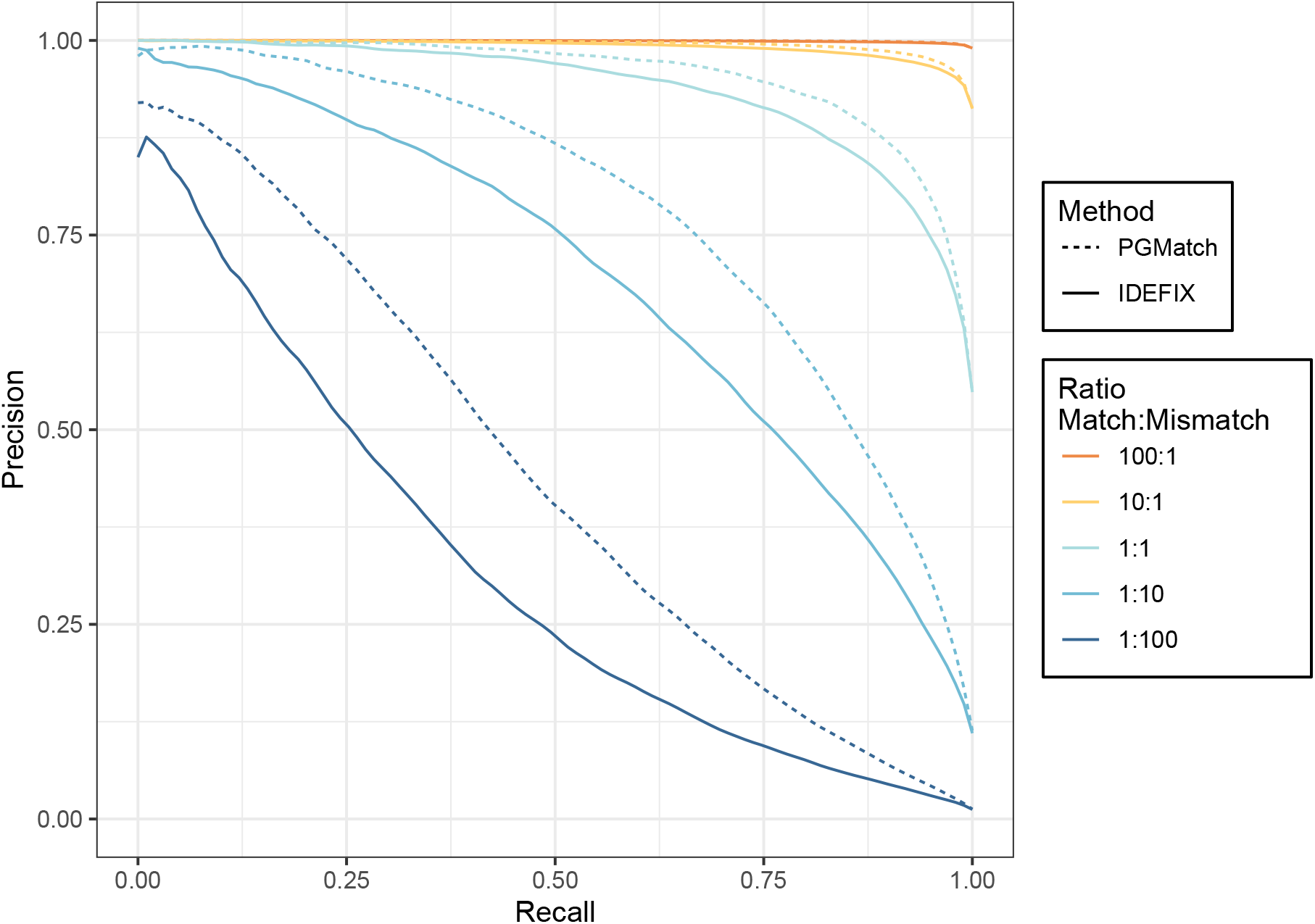
Comparison with an existing method. Precision–recall curves for PGMatch (dashed) and IDEFIX (solid) under varying match-to-mismatch ratios. Warmer colors indicate higher proportions of matches relative to mismatches.

### Inference of sensitive genomic variants

The re-identification of a genome creates a privacy risk because of the sensitive information a genome could disclose about his owner. For example, one could check the APOE status of an individual through his genome to infer his risk of developing Alzheimer’s disease, as the APOE-*ε*4 haplotype is a risk factor for this disease. In the previous sections, we showed that re-identification by phenotypic prediction cannot reliably find out who is the owner of an anonymized genome. However, one could argue that the individuals whose PGS matches the focal sample’s phenotypes with a high probability might be genetically similar to the target individual, and thus might also share sensitive genetic variants with him. For example, one can guess the probability that an individual *I* with a known phenotype profile carries a genetic variant of interest “V” (for instance APOE-*ε*4) based on the following estimate:

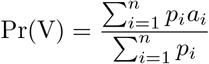

where *p*_*i*_ is the match-probability between the target individual, *I*, and biobank sample *i*, and *a*_*i*_ is a 0-1 indicator variable whether the genome of individual *i* carries the variant of interest.

To assess the accuracy of this approach, we evaluated multiple genetic variants associated with diverse traits and diseases, including APOE-*ε*4 (Alzheimer’s disease risk), *ABO* (blood group), *LCT* (lactose tolerance), *FUT2* (secretor status), and *HLA-DQB1*06:02* (narcolepsy risk). For each genetic variant, we computed Pr(V) for 1 000 target individuals based on comparisons to 100 000 genomes. We first compared the distributions of predicted carrier probabilities between true carriers and non-carriers (APOE in **Figure 6A,B**, other genetic variants in **Supp. Fig. S8-11 A,B**). A significant difference between these distributions was observed for APOE-*ε*4 and *ABO*, whereas no significant difference was observed for *FUT2* and *HLA-DQB1*06:02*. For *LCT*, the difference between carriers and non-carriers was only marginally significant. We then repeated the analysis over 100 bootstrap replicates and evaluated classification performance using precision–recall curves (APOE in **Figure 6C,D**, other genetic variants in **Supp. Fig. S8-11 C,D**). For genetic variants showing no significant separation between carriers and non-carriers (*FUT2* and *HLA-DQB1*06:02*), precision remained close to the population frequency across all recall levels, indicating no meaningful enrichment of carriers among high-probability individuals. Similarly, for *LCT*, the weak separation between distributions resulted in little to no improvement in precision as recall decreased. In contrast, for genetic variants with significant differences in predicted probabilities (APOE-*ε*4 and *ABO*), precision increased as recall decreased, indicating enrichment of carriers among individuals with the highest predicted probabilities. Thus, among individuals with the top 5% of the highest predicted probability of being carriers, the mean variant carrier frequency across the 100 bootstrap replicates was markedly higher (0.56 for APOE-*ε*4 and 0.60 for *ABO*) than these frequencies in the overall population (0.29 and 0.44, respectively).

**Figure 6:**
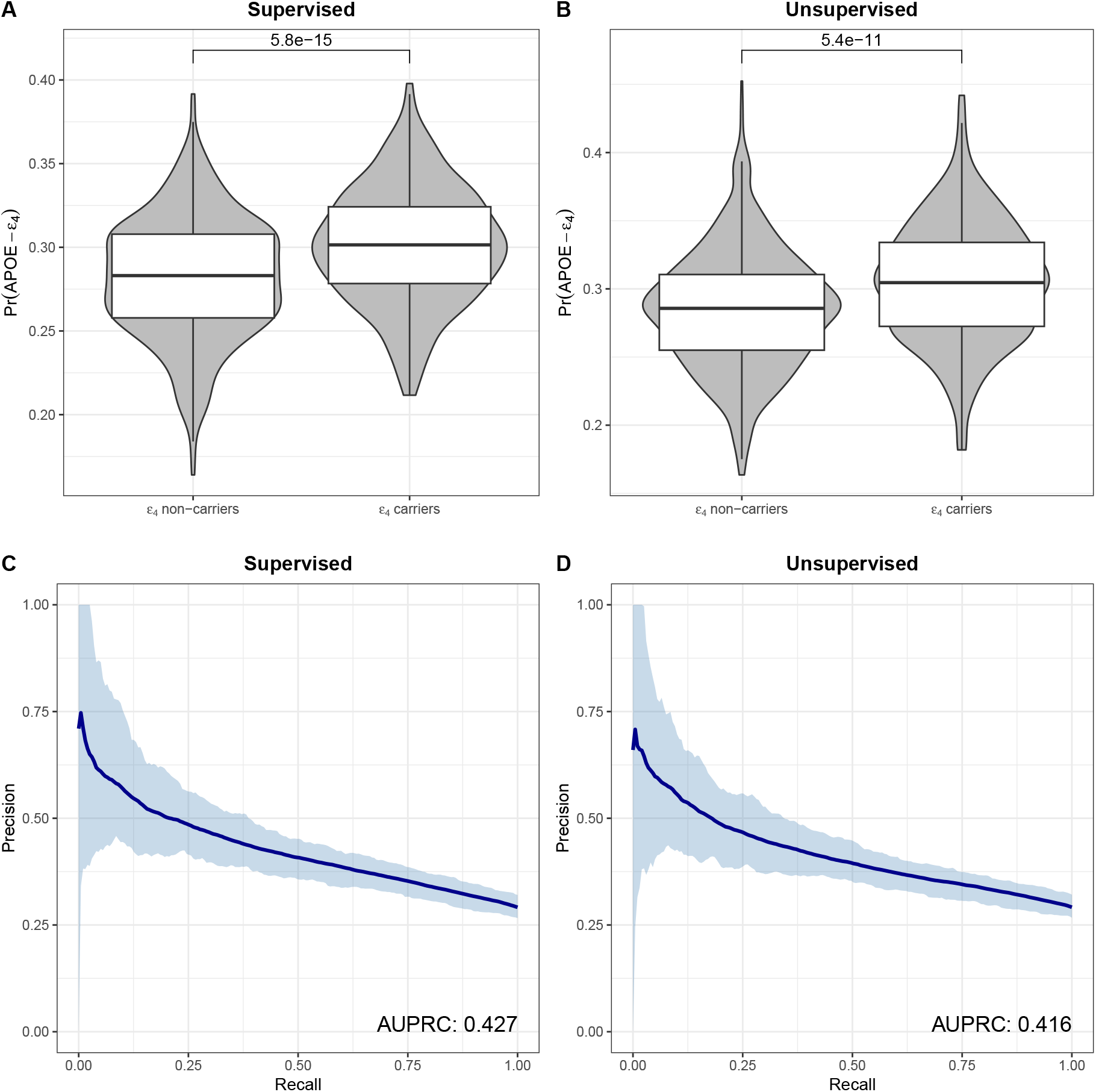
Inference of APOE-*ε*4 status using re-identification by phenotypic prediction. **A, B**. Distribution of Pr(APOE-*ε*4) in *ε*4 carriers and non-carriers using the supervised and unsupervised approaches, respectively. Results of one-sided *t*-tests are shown above the connecting bars. **C, D**. Precision-recall curves for the inference of *ε*4 carriers across different Pr(APOE-*ε*4) thresholds using the supervised and unsupervised approaches, respectively. The thick blue line represents the mean precision at a given recall level across 100 experiments. Ribbons represent the 95% bootstrap percentile interval of precision across experiments.

To explain differences in prediction accuracy across genetic variants, we hypothesized that they primarily reflect differences in their association with the phenotypes used for re-identification. To test this, we randomly sampled 10 000 individuals per genetic variant, trained a generalized linear model (GLM) on 1 000 individuals, and predicted variant status as a function of the phenotypes in the remaining 9 000, repeating this procedure over 100 bootstraps **Supp. Fig. S12**. Precision-recall curves closely matched those obtained via re-identification, indicating that sensitive genetic variant inference is largely driven by variant–phenotype correlations. Thus, access to genomic data provides little additional information beyond what is already captured by phenotypes.

## Discussion

In this work, we analyzed the risk of re-identification by phenotypic prediction. We developed a state-of-the-art method tailored for this aim, relying on PGS while taking into account genetic and environmental correlations between phenotypes: PGMatch. We compared our approach to IDEFIX, a method relying on the comparison of PGS and phenotypes for detecting sample mismatches within biobanks. However, IDEFIX is conceptually different and cannot address the scenario considered under our unsupervised approach, which more closely reflects real-world conditions. In this setting, our method requires only summary statistics, along with the genomes and phenotypes of the target individuals, whereas IDEFIX requires a large dataset to learn the relationship between PGS and phenotypes. Nevertheless, IDEFIX represents the closest methodological analog currently available, and we, therefore, used it as a benchmark for comparison with our supervised approach. Our method demonstrated a higher computational efficiency and predictive performance. The lighter memory usage and running time of our approach mainly result from the way the existing tool handles the computation of the *LLR*: For a dataset of *n* individuals, the former method computes and stores the *LLR* between all possible phenotype–PGS combinations, i.e., *n ×n LLR* values, while we compute the *LLR* only between *n* phenotypes and *n* PGSs. This performance advantage renders re-identification by phenotypic prediction between an individual and an entire biobank of hundreds of thousands of genomes feasible, even on a personal computer. Regarding predictive performance, the difference is due to the following two reasons. First, while our approach weights the PGS-phenotype match likelihoods based on their expected similarity, the reference method does not. Instead, it filters phenotypes based on a t-test comparing the residuals of matches to those of mismatches. Thus, the discrepancy between a trait and a PGS for a phenotype that is easily predictable (e.g. standing height) is considered equally important in the re-identification as the trait-predictor deviation for a phenotype that is harder to predict (e.g. education age), increasing the overlap of the *LLR* distribution under *H*_0_ with the *LLR* distribution under *H*_1_. Second, the baseline method does not account for the correlations between the phenotypes, which can lead to an overestimation of the difference between these two distributions. As an example, in the extreme case where the same phenotype (e.g. standing height) is mistakenly counted twice, this method would estimate a stronger distinction between the *LLR* of matches and mismatches, even though these two instances of the same phenotype contain the same information.

These two factors also explain the non-linear improvement of our method as the number of phenotypes increases. In our experiments, we sorted the phenotypes according to the proportion of variance explained by genetics, and the results reported in **Figure 2,C–F** correspond to the top 10, 20, 30, and 40 phenotypes with the highest explained variance. This means that the phenotypes added between 10 and 20 are more predictable and therefore contribute more to re-identification than those added between 30 and 40. Moreover, the newly added phenotypes may be partially correlated with phenotypes already included, which limits their contribution (even though we filtered out phenotypes with a correlation *>* than 0.8). Interestingly, our method performed better in the inference of matches than mismatches). In both supervised and unsupervised approaches, this can be explained by the way the *LLR* distributions under *H*_0_ and *H*_1_ overlap **Supp. Fig. S1**; the difference in density between *H*_0_ and *H*_1_ is more pronounced at high *LLR* values (corresponding to matches) than at low *LLR* values (corresponding to mismatches. In **Supp. Fig. S1**, we also notice that many individuals’ phenotypes match poorly with the predicted traits based on their own genome (extremely low *LLR* values), which affects the performance of the unsupervised approach more than that of the supervised counterpart. This strong deviation between an individual’s predicted trait and actual trait can be explained by other factors not captured by the PGS but having an important effect on the phenotype. For instance, Hawkes et al. showed that UK Biobank participants whose observed height deviated strongly from their genetically predicted height were enriched for congenital malformations and carried loss-of-function variants in genes associated with growth disorders [14].Such cases illustrate that individuals affected by rare genetic diseases may be more difficult to re-identify if the disease status is unknown, since their phenotypes deviate from polygenic expectations.

Despite the strong performance of our method on a balanced dataset of matches and mismatches, its precision dropped drastically when assessed in more realistic scenarios, where encountering a non-matching individual–genome pair is more likely than a matching one (**Figure 4A-B** and **Table 2**). Using the UK Biobank as an example, we showed that even if one knew that the individual was part of the biobank (prior (C)), the AUPRC would remain below 4 *×*10^−3^. These results highlight the difficulty of re-identification via phenotypic prediction in real-world settings. More generally, to assess the re-identification risk of a genome based on its predicted phenotypes, one should consider two key factors: the prior probability of a match and the prediction accuracy of the PGSs for the respective phenotypes.

A high prior probability may arise in constrained search spaces, such as small or highly structured cohorts (e.g., isolated populations or narrowly defined clinical studies), where the number of plausible genome–individual pairings is limited. For example, in datasets such as the Fenland Study (12 435 participants) or the 1000 Genomes Project (2 504 participants), knowing that an individual is included in the dataset would lead to higher AUPRCs than in the UK Biobank [15, 16].Access to auxiliary information, such as ancestry, age range, or geographic location, can further restrict the pool of candidate individuals, thereby increasing the prior probability of a match. For instance, only 105 Finnish individuals are included in the 1000 Genomes Project; knowing that the target individual is Finnish and part of the dataset would raise the prior to approximately 1/100. However, as shown in **Figure 4**, even under these conditions the AUPRC reaches only 0.4416, indicating that re-identification via polygenic prediction remains highly unlikely in typical biobank settings. In the context of genome re-identification within a biobank, where every genome is equally likely to belong to the target individual, the prior probability cannot exceed 0.5, as higher values would imply that only a single genome is present, making the match trivial. Nevertheless, there are scenarios in which the prior probability of a match may exceed 0.5. One example is erroneous biological sampling. A related case is addressed by IDEFIX, which focuses on detecting sample mislabeling in biobanks using genetic sex versus self-reported sex as a prior. In a validation setting (not considered in their study), the prior probability of a correct match would be substantially greater than 0.5, as sample mix-ups are expected to be rare in biobanks. Another context in which the prior can be greater than 0.5 is forensic analysis. In such settings, additional contextual information from a crime scene can increase the prior probability that a given DNA sample originates from a specific individual. Relatedly, forensic DNA phenotyping (FDP) is increasingly used to infer externally visible characteristics from DNA and support investigative hypotheses about an unknown individual [17].FDP currently relies on a limited set of phenotypes predicted from a small number of genetic variants, which underscores the gap between the theoretical best-case scenarios considered in this work and current real-world practice. Another context in which the prior might be higher than 0.5 is parental testing. Let *G* be the genome of a child and *I* be a potential parent with multiple known phenotypes. If the parent-of-origin of each haplotype could be inferred [18, 19], a polygenic score (PGS) could be computed on *G* for the corresponding parental haplotypes and then compared to the phenotypes of *I*, further increasing match identification. However, this approach goes beyond the scope of re-identification by phenotypic prediction and would require a dedicated study to address the associated challenges and assess its practical feasibility. To explore regimes with priors exceeding 0.5, we included scenarios with 100:1 and 10:1 match:mismatch ratios in our analysis of prior effects (**Figure 4**, top two warmest colors). These results show that in such contexts re-identification can increase the probability of a correct match beyond the original prior. An adversary using PGMatch could adapt the method to take a different prior than 1:1 and recompute the posterior accordingly.

On the other hand, prediction accuracy could be further improved by incorporating additional sources of information. In particular, rare genetic variants may provide complementary signal to common variants, especially for traits influenced by low-frequency but high-impact alleles. Individuals carrying such variants may also exhibit extreme phenotypes, thereby reducing the number of potential matches and increasing re-identification risk. However, the impact of rare variants is highly disease-specific and would require a dedicated study to be properly assessed. We therefore acknowledge that individuals with rare genetic conditions may be at increased risk of re-identification, but a systematic evaluation of this effect is beyond the scope of the present work. Second, the inclusion of additional phenotypic data could substantially enhance predictive performance. Any covariate that partially explains the phenotypes of interest would improve prediction accuracy. For instance, routinely collected information in electronic health records (EHRs)—such as clinical measurements, diagnoses, medication history, and longitudinal health trajectories—could serve as informative covariates to refine phenotype prediction. Thus, integrating both richer genetic variation and more detailed phenotypic descriptors could improve the overall accuracy of re-identification models beyond what is achievable using common variants and limited phenotype sets alone. However, regardless of improvements in phenotypic prediction, re-identification remains fundamentally constrained by the extremely low prior probability of a match in realistic settings, as mentioned above. To better assess the difficulty of such re-identification, the adversary can estimate the expected success probability from the *LLR* distributions derived within our framework. In essence, successful identification of a match depends on the estimated *LLR* distribution under *H*_1_, which determines the probability that the match score exceeds a chosen decision threshold. While this approximation may be less accurate for mismatch inference under the unsupervised approach, as discussed above, it should provide a reasonable estimate for match inference in both models.

Apart from the aforementioned limitations, the main factor making re-identification by phenotypic prediction unlikely, as already noted by Erlich in 2014 [6], is the availability of the required datasets. Access to genetic data remains tightly regulated in many countries, and even when pseudonymized, such data are still considered personal information under the GDPR, justifying their restricted access [20].Furthermore, even if an attacker was able to bypass regulations and obtain an anonymized genome, they would also need access to highly predictable traits of the target individual. To our knowledge, no dataset currently exists that contains both such phenotypes and the corresponding identities, and any such resource would fall under GDPR regulations on personal data.

Another key limitation of this approach lies in the challenges posed by ancestry in phenotype prediction. In our study, we did not explicitly model genetic ancestry in the computation of polygenic scores. Instead, all analyses were restricted to individuals of White British ancestry in the UK Biobank (as defined by field 22006 from UK Biobank), thereby considering a relatively homogeneous population. This design reflects a best-case scenario for re-identification, as polygenic scores are known to achieve their highest predictive performance within ancestrally matched populations. In more heterogeneous settings, the portability of polygenic predictors is substantially reduced, leading to lower prediction accuracy and consequently reduced re-identification performance. Therefore, our results likely represent an upper bound on the effectiveness of phenotype-based re-identification. In practice, the presence of diverse ancestries within a dataset would further decrease the feasibility of such approaches, reinforcing the conclusion that re-identification remains challenging under realistic conditions.

Methods have also been developed to mitigate re-identification by phenotypic prediction in genetic datasets. The challenge in this case lies in modifying genetic data while preserving their research utility. For instance, approaches have been proposed to make genetic datasets resistant to re-identification by phenotypic prediction while maintaining key properties required for them to act as reference panels in imputation contexts [21, 22, 23].These methods often involve shuffling haplotype segments, which disrupts genome–phenome links and renders re-identification by phenotypic prediction impossible. Another approach consists of rendering genetic data completely anonymous yet still analysable, for example through homomorphic encryption [24, 25].

Finally, we assessed whether re-identification by phenotypic prediction could be used to perform genetic variants inference, that is, to infer whether an individual was carrying a specific variant of interest. Among the tested variants, the highest predictive performance was reached for APOE-*ε*4, a known risk factor of Alzheimer’s disease [12, 26].However, this approach was not very efficient, as if an attacker claimed people to be APOE-*ε*4 carriers whenever Pr(APOE-*ε*4) *>* 0.330, they would miss 80% of the true APOE-*ε*4 carriers, and half of their claimed carriers would actually not carry the APOE-*ε*4 variant (**Figure 6**). More importantly, most of the predictive power was due to the correlation between the variants and the phenotypes used for re-identification. Thus, access to genomic data provides little additional information beyond what is already captured by phenotypes. These conclusions were consistant across the four other variants we tested. Moreover, it is important to note that APOE-*ε*4 is only a risk factor; carrying it does not imply certain development of Alzheimer’s disease. For instance, among individuals above 80 years old, even those with two copies of the APOE-*ε*4 variant have at most a 24% chance of developing Alzheimer’s disease [26].This example highlights the actual threat raised by knowing one’s genome: it can be used to infer sensitive phenotypes which may be used for discrimination. However, the precision of the inference strongly varies depending on the phenotype, its heritability and the availability of relevant summary statistics. Phenotype inference through a genome remains a very challenging task, which not only makes re-identification of a genome difficult, but also limits the extent of what can be inferred from a re-identified genome.

## Materials and Methods

### Method description

Based on *T* traits of an individual *I* and the corresponding *T* PGSs of a genome *G*, our method aims to distinguish between two scenarios: *G* belongs to *I* (*H*_1_) and *G* does not belong to *I* (*H*_0_) (**Figure 1**). The method is divided into two steps: first, it computes a log-likelihood ratio (*LLR*) of observing the traits and PGSs under *H*_0_ *vs H*_1_, and then it transforms this *LLR* into a probability by comparing it to the expected *LLR* distribution under *H*_0_ and *H*_1_.

### LLR calculation

Let the *T* traits of the individual be denoted as *y* = (*y*_1_, *y*_2_, …, *y*_*T*_) and the *T* corresponding PGSs (computed from the genome) be denoted as *s* = (*s*_1_, *s*_2_, …, *s*_*T*_). Let us assume that we know the proportion of phenotypic variance explained by the PGSs for each of the *T* traits 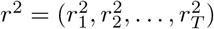 (**Figure 1,A**). Each trait can be decomposed into a genetic component (approximated by the polygenic score) and a residual component and hence be expressed as:

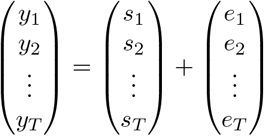

or in vector form **y** = **s** + **e** with

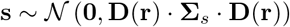

And

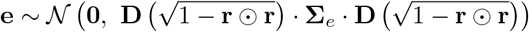

where Σ_*s*_ and Σ_*e*_ represent the correlation between the polygenic scores and the correlation between the residuals, respectively, and *D*(*v*) denotes a diagonal matrix with diagonal elements defined by vector *v*. In other words, ***s*** and ***e*** follow multivariate normal distributions with correlation matrices Σ_*s*_ and Σ_*e*_, respectively, and variances *r*^2^ and 1 *− r*^2^ for each component of the distributions.

Let 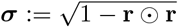, i.e. the vector of standard deviations of the non-genetic (residual) components for each trait. To compare the likelihood of observing ***y*** and ***s*** under *H*_0_ and *H*_1_, we compute their log-likelihood ratio as:

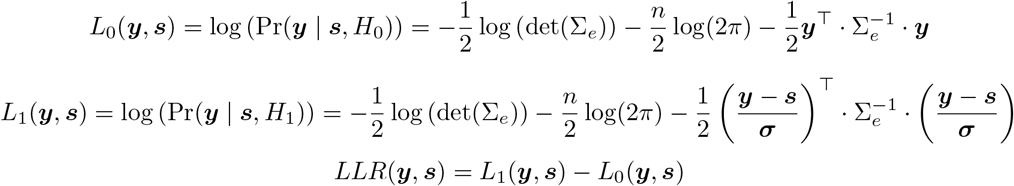

We expect *LLR*(***y, s***) being larger when ***y*** and ***s*** come from the same person than when they come from different people.

### LLR transformation into probability

In order to transform *LLR*(***y, s***) into a probability of ***y*** and ***s*** coming from the same individual (Pr(*G* = *I* | *LLR*)), we need to estimate the expected *LLR* distribution under *H*_0_ and *H*_1_ (**Figure 1B**). The way we estimate these distributions depends on whether we have access only to association summary statistics (**unsupervised approach**) or we also have access to a dataset of samples with genomes and phenotypes (**supervised approach**).

#### Supervised approach

In this situation, we simply compute *LLRs* in the available dataset between all individuals and their respective genomes (*H*_1_), and between all individuals and randomly sampled genomes (*H*_0_). To do so, we derive Σ_*e*_, Σ_*s*_ and *r*^2^ directly from the training dataset. We describe how we obtain Σ_*e*_,Σ_*s*_ and *r*^2^ in the **Data** subsection. From the obtained *LLR* values, we estimate the expected distribution of the *LLRs* under *H*_0_ and *H*_1_ using kernel density estimation.

#### Unsupervised approach

Here, we need to simulate a dataset of phenotypes and PGSs. As described earlier,

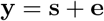

which means

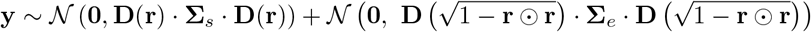

We can simulate *s* and *y* solely using Σ_*e*_,Σ_*s*_ and *r*^2^, which are derived from summary statistics in this case (see corresponding **Data** subsection). In practice, we simulate 500 000 pairs of individuals and PGSs and we repeat the process aforementioned in the **Supervised approach**. In this case, the expected distribution of the *LLRs* under *H*_0_ and *H*_1_ are estimated on simulated values, again using kernel density estimation.

Let 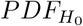 and 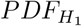 be the probability density functions of the approximated *LLRs* distribution (estimated through kernel density smoothing) under *H*_0_ and *H*_1_, respectively. For a new pair of individual and genome (**Figure 1C**), we can compute the probability of *y* and *s* coming from the same individual (a match) as

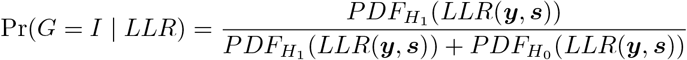

To compute the probability that ***y*** and ***s*** come from two different individuals (a mismatch), we simply use the expression

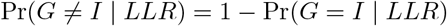

### Probability of being part of a Biobank

To compute the number of individuals leaving in a 40km radius of the assessment centres, we used data from the Office of National Statistics. They provide population estimates per year for Lower layer Super Output Area population (LSOA), which corresponds to small geographic areas used by England and Wales for statistics. For each assessment centre, we selected the LSOA in a radius of 40km from them using their latitude and longitude coordinates and the Haversine formula. Then we counted the number of people between 40 and 74 years of age in 2011 in these areas (*T*_*LSOA*_) that we downloaded from here.Some of the LSOA were absent from the statistics of 2011, which means the estimate we report in **Supp. Tab. S2** might be slightly lower than the real count.

Next, we retrieved the total number of UK Biobank participants recruited in the respective assessment centre *T*_*assessment*_. We accessed these data through the field “Locations of UK Biobank assessment centres throughout United Kingdom” from the UK Biobank accessible here.Finally, we computed the probability of a randomly sampled genome from the UK Biobank to be owned by a randomly sampled individual from a specific area as 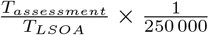.

Other methods could be used to narrow down the number of genomes potentially matching an individual. For instance, one could perform principal component analysis on the genomes of the UK Biobank to infer if they were more likely to come from a specific assessment centre. However, such approaches are dataset-specific, and fall out of the scope of this paper. Here, we focus in the effect of the *a priori* probability that a genome matches an individual, the consequence of which we studied in the **Results** section.

### Data

In this subsection, we describe the summary statistics we used in our analysis. Also, we describe how we obtained the GWAS summary statistics required to compute the PGSs, the correlation between PGSs Σ_*s*_, the environmental correlation Σ_*e*_ and the *r*^2^ for each phenotype. We also explain how we treated the phenotypes in our experiments and defined carrier status in the inference of sensitive genomic variants.

#### Summary statistics

We downsampled the UK Biobank into a training set of randomly selected 150 000 white British individuals and performed GWASs on them for 76 traits. In our analysis, we only kept the phenotypes (i) present in *>* 100 000 individuals of each sex and (ii) with a correlation *<* 0.8 (list of 59 resulting traits studied available in **Table S1**). To perform the GWASs, we used REGENIE [27]with parameter *–bsize 1000* in step1 and parameters *–bsize 400 –firth –approx –pThresh 0*.*01* in step2. We used 10 Principal Components, age and sex as covariates as previoulsy described [19].We then adjusted the summary statistics based on LD using LDpred2 (bigsnpr v 1.12.18) [28].As advised in the tutorial, we restricted our analysis to the HapMap3+ variants and used the corresponding LD matrices covering these SNPs. We used the *snp ldsc()* function to estimate heritability and provided these estimates to the *snp ldpred2 auto()* function as starting point with the following parameters: *burn in=500, num iter=500, report step=20, allow jump sign=F, use MLE=T, shrink corrr=0*.*95, vec p init=seq log(1e-4, 0*.*2, length*.*out = 10)*.

#### Polygenic scores

We used the resulting summary statistics to compute the PGSs of the white British individuals not used for the GWASs (409 599 - 150 000 = 259 599 individuals). Briefly, we calculated the PGSs by computing the weighted sum of allele dosages across genetic variants without adjustment for additional covariates, as we assume the adversary aiming at re-identifying the target individual has access only to genetic data and no auxiliary information. To do so, we used the *pgs calc* software [29].The resulting variance explained by the PGSs in each phenotype can be found in **Table S1**.

#### Model parameter estimation

In the **supervised approach**, the environmental correlation matrix Σ_*e*_ is estimated by computing the Pearson correlation between phenotypes in the training set. The PGS correlation matrix Σ_*s*_ is estimated by computing the residuals between phenotypes and PGSs in the training set and calculating the correlation matrix of the resulting values. For each phenotype, *r*^2^ is computed as the squared Pearson correlation between *y* and *s* in the training set.

In the **unsupervised approach**, Σ_*e*_ is assumed to be equal to Σ_*s*_, as environmental correlations are not accessible without individual-level data. The PGS correlation matrix Σ_*s*_ is approximated using the genetic correlation matrix Σ_*g*_ estimated from GWAS summary statistics using LD score regression with the *ldsc()* function from GenomicSEM (v. 0.0.5) [30].Similarly, *r*^2^ is approximated using trait heritability estimated with the same software.

#### Phenotype and PGS pre-processing

In all experiments, the phenotypes were normalized and corrected for sex. The sex correction allowed us to increase the number of comparisons we could make, as males and females could then be compared. To improve phenotype prediction, we could have also corrected for age or other covariates. However, in this study, we assumed that the attacker had access only to phenotypes that can be predicted from genetic data.

The PGSs were normalized using the mean and standard deviation of individuals in the generated datasets. In practice, an adversary with access to only a single genome would need to derive the mean and standard deviation of the PGS in the population from summary statistics alone.

The expected population mean PGS can be computed analytically as:

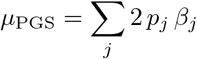

and the population variance of the PGS as:

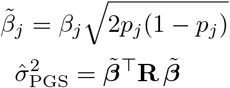

where *p*_*j*_ is the effect allele’s frequency at locus *j, β*_*j*_ is its effect size, and **R** is the population LD matrix. Both quantities are thus fully recoverable from GWAS summary statistics and a reference LD panel, without requiring access to individual-level data.

## Definition of carrier status

Carrier status was defined as follow: For APOE, carrier status was defined as the presence of at least one APOE-*ε*4 haplotype, determined from the combination of rs429358 and rs7412 genotypes. For *LCT* and *FUT2*, individuals were classified as carriers if they possessed at least one effect allele at rs4988235 and rs601338, respectively. For *ABO*, carrier status was defined as homozygosity for the O haplotype. The O haplotype was determined following the criteria used in UK Biobank data-field 2315: “*rs8176719 deletion was taken as indicative of haplotype O; For participants with no result for rs8176719, rs505922 of T was used to indicate type O*.”. For *HLA-DQB1*06:02*, carrier status was determined using imputed haplotype calls (data-field 22182).

## Data availability

This research was conducted using phenotype and genotype and imputed data from the UK Biobank. The UK Biobank data are available under restricted access, which can be obtained by application via the UK Biobank Access Management System (https://www.ukbiobank.ac.uk/enable-your-research/apply-for-access. In addition, we used statistics from the UK Office of National Statistics, available here.

## Code availability

All source code an analysis scripts are publicly available at https://github.com/TheoCavinato/PGMatch. The following tools were used in our analysis:

- REGENIE (https://github.com/rgcgithub/regenie)
- GenomicSEM (https://github.com/GenomicSEM/GenomicSEM)

## Author contributions

Z.K. and T.C. conceptualized the study and wrote the manuscript. Z.K. designed the statistical tests. T.C. developed the algorithms and performed all experiments. R.J.H. generated the GWAS summary statistics. All authors contributed to the interpretation of the results, revised the manuscript, and approved the final version. The project was supervised by Z.K.

## Acknowledgements

This work has been funded by the Swiss National Science Foundation (315230-219587) and conducted using the UK Biobank Resource under Application Number 16389. The UK Biobank data analysis was conducted using the High-Performance Computing Center at the University of Lausanne, Switzerland. T.C. was funded by the Department of Computational Biology, Lausanne, Switzerland.

