## Supplements for "Evaluating anonymized genome re-identification using polygenic predictions and its implications for data privacy"

### Supplementary Notes

#### *Supplementary Note 1 - Assessing if the target is part of the biobank*

To assess whether an individual belongs to a biobank, one approach would consist of applying our method to compare the individual with each genome in the biobank, sum the resulting matching probabilities and calculate the mean as:

$$\frac{1}{n}\sum_{i=1}^{n} p_i$$

where $p_i$ is the match-probability between the target individual, $I$, and biobank sample $i$ in a biobank of size $n$. If this mean is larger when the individual is part of the biobank than when they are not, it could provide means to discriminate between the two scenarios. To test this idea, we randomly selected 1 000 individuals, compared them to 100 000 genomes (including/ or not their own genome), and calculated the resulting means of the matching probabilities. Unfortunately, the difference in the median of the two scenarios was only of $-8.2 \times 10^{-6}$, preventing reliable discrimination (see distributions in **Supp. Figure S13 A**).

One could also argue that, if an individual is part of the biobank, the maximum matching probability when comparing the individual with each genome would be higher than if he was not. But again, the difference was small ($1.4 \times 10^{-7}$) between the medians of the two scenarios (**Supp. Figure S13 B**).

For both metrics, the difference between the two scenarios would increase as the biobank size decreases. For example, in the extreme case of a "biobank" containing only two individuals, the metrics would clearly distinguish whether the target individual is included. However, contemporary biobanks typically comprise more than 100 000 participants, rendering this approach ineffective in realistic settings.

### Supplementary Tables and Figures

**Table S1.** Phenotypes used in this study, their corresponding UKB identifier and their sex-adjusted proportion of phenotypic variance explained by their PGSs.

| Phenotype name | UKB identifier | $r^2$ |
|---|---|---|
| Mean platelet (thrombocyte) volume | 30100 | 0.3529248 |
| Standing height | 50 | 0.3279128 |
| Hair colour (natural, before greying) | 1747 | 0.2788137 |

| | | |
|---|---|---|
| Total bilirubin (umol/L) | 30840 | 0.2584552 |
| Platelet count | 30080 | 0.2166321 |
| Mean corpuscular haemoglobin | 30050 | 0.2160767 |
| Platelet distribution width | 30110 | 0.2069917 |
| Alkaline phosphatase (U/L) | 30610 | 0.1899249 |
| HDL cholesterol (quantile) | 30760 | 0.1854733 |
| Monocyte count | 30130 | 0.1600537 |
| Mean reticulocyte volume | 30260 | 0.1587511 |
| SHBG (quantile) | 30830 | 0.157124 |
| Ease of skin tanning | 1727 | 0.1563104 |
| Mean sphered cell volume | 30270 | 0.1527701 |
| Whole body water mass | 23102 | 0.1516035 |
| Red blood cell (erythrocyte) distribution width | 30070 | 0.1463586 |
| Red blood cell (erythrocyte) count | 30010 | 0.1405535 |
| Gamma glutamyltransferase (U/L) | 30730 | 0.1402482 |
| Apoliprotein B (quantile) | 30640 | 0.1397056 |
| Cystatin C (quantile) | 30720 | 0.1389114 |
| Urate (umol/L) | 30880 | 0.1372439 |
| Glycated haemoglobin (quantile) | 30750 | 0.1370882 |
| High light scatter reticulocyte count | 30300 | 0.1363926 |
| Eosinophill count | 30150 | 0.1287876 |
| IGF-1 (quantile) | 30770 | 0.1261221 |
| Triglycerides (quantile) | 30870 | 0.121383 |
| C-reactive protein (quantile) | 30710 | 0.1196107 |
| Lymphocyte count | 30120 | 0.1114969 |
| Creatinine (umol/L) | 30700 | 0.1073002 |
| White blood cell (leukocyte) count | 30000 | 0.1003281 |
| Body mass index (BMI) | 21001 | 0.1003243 |
| Immature reticulocyte fraction | 30280 | 0.08764611 |
| Total protein (quantile) | 30860 | 0.08455383 |
| Aspartate aminotransferase (U/L) | 30650 | 0.07668835 |
| Forced vital capacity (FVC) | 3062 | 0.06983692 |
| Calcium (quantile) | 30680 | 0.06424278 |
| Albumin (quantile) | 30600 | 0.06317011 |
| Education age | 8451 | 0.05891298 |
| Phosphate (quantile) | 30810 | 0.05466916 |
| Alanine aminotransferase (U/L) | 30620 | 0.05391909 |
| Vitamin D (quantile) | 30890 | 0.04737091 |
| Urea (quantile) | 30670 | 0.04559821 |
| Diastolic blood pressure, automated reading | 4079 | 0.04529444 |
| Systolic blood pressure | 4080 | 0.04353869 |
| Glucose (quantile) | 30740 | 0.03669467 |
| Chronotype | 11801 | 0.02780183 |
| Testosterone (quantile) | 30850 | 0.02749741 |
| Time spent watching TV | 1070 | 0.02375606 |
| Ever Smoked | 20116 | 0.02112763 |
| Neuroticism score | 20127 | 0.02065047 |
| Alcohol Consumption (Frequency) | 1558 | 0.01708822 |
| Overall health rating | 2178 | 0.01635662 |
| Type 2 diabetes | E11 | 0.01590447 |
| Basophill count | 30160 | 0.01554728 |
| Educational attainment | as described in [1] | 0.01126346 |
| Sleep duration | 1160 | 0.01107786 |
| Walking pace | 924 | 0.008047851 |

| | | |
|---|---|---|
| Number of Years of Education | 6138 | 0.002593543 |
| Nucleated red blood cell count | 30170 | 2.499677e-07 |

**Table S2.** Assessing the probability for a randomly sampled individual leaving in a 40km radius from the assessment centre and matching UKB's recruitment criteria to match with a randomly sampled genome from the UKB.

| Assessment centre | Participants in assessment centre | Total number of individuals in a 40km radius from assessment centre | Match:Mismatch ratio |
|---|---|---|---|
| Birmingham | 25'506 | 1'781'250 | 1:17'459'127 |
| Bristol | 43'020 | 841'131 | 1:4'888'023 |
| Cardiff | 17'885 | 828'030 | 1:11'574'364 |
| Cheadle | 20'348 | 1'877'998 | 1:23'073'496 |
| Croydon | 27'392 | 3'967'821 | 1:36'213'320 |
| Hounslow | 28'881 | 4'125'233 | 1:35'708'883 |
| Leeds | 44'220 | 1'353'097 | 1:7'649'802 |
| Liverpool | 32'825 | 1'429'777 | 1:10'889'391 |
| London Barts | 12'584 | 4'088'077 | 1:81'215'770 |
| Manchester | 13'943 | 2'049'482 | 1:36'747'508 |
| Middlesbrough | 21'290 | 621'147 | 1:7'293'882 |
| Newcastle | 37'011 | 847'571 | 1:5'725'129 |
| Nottingham | 33'883 | 1'186'263 | 1:8'752'641 |
| Oxford | 14'063 | 545'854 | 1:9'703'726 |
| Reading | 29'426 | 1'159'326 | 1:9'849'504 |
| Sheffield | 30'399 | 1'340'404 | 1:11'023'422 |
| Stockport | 3'799 | 1'872'866 | 1:123'247'302 |
| Stoke | 19'441 | 702'657 | 1:9'035'762 |
| Swansea | 2'284 | 386'646 | 1:42'321'147 |
| Wrexham | 649 | 664'166 | 1:255'842'065 |

**Figure S1. Effect of the number of phenotypes on LLR distributions under the supervised approach and the unsupervised approach across different number of phenotypes.** Distribution of the *LLRs* values expected from summary statistics or a train set (solid lines) and observed in a test set (dashed lines) when comparing a genome to its actual traits ($H_1$ in green) and to the traits of a randomly sampled other individual ($H_0$ in red).

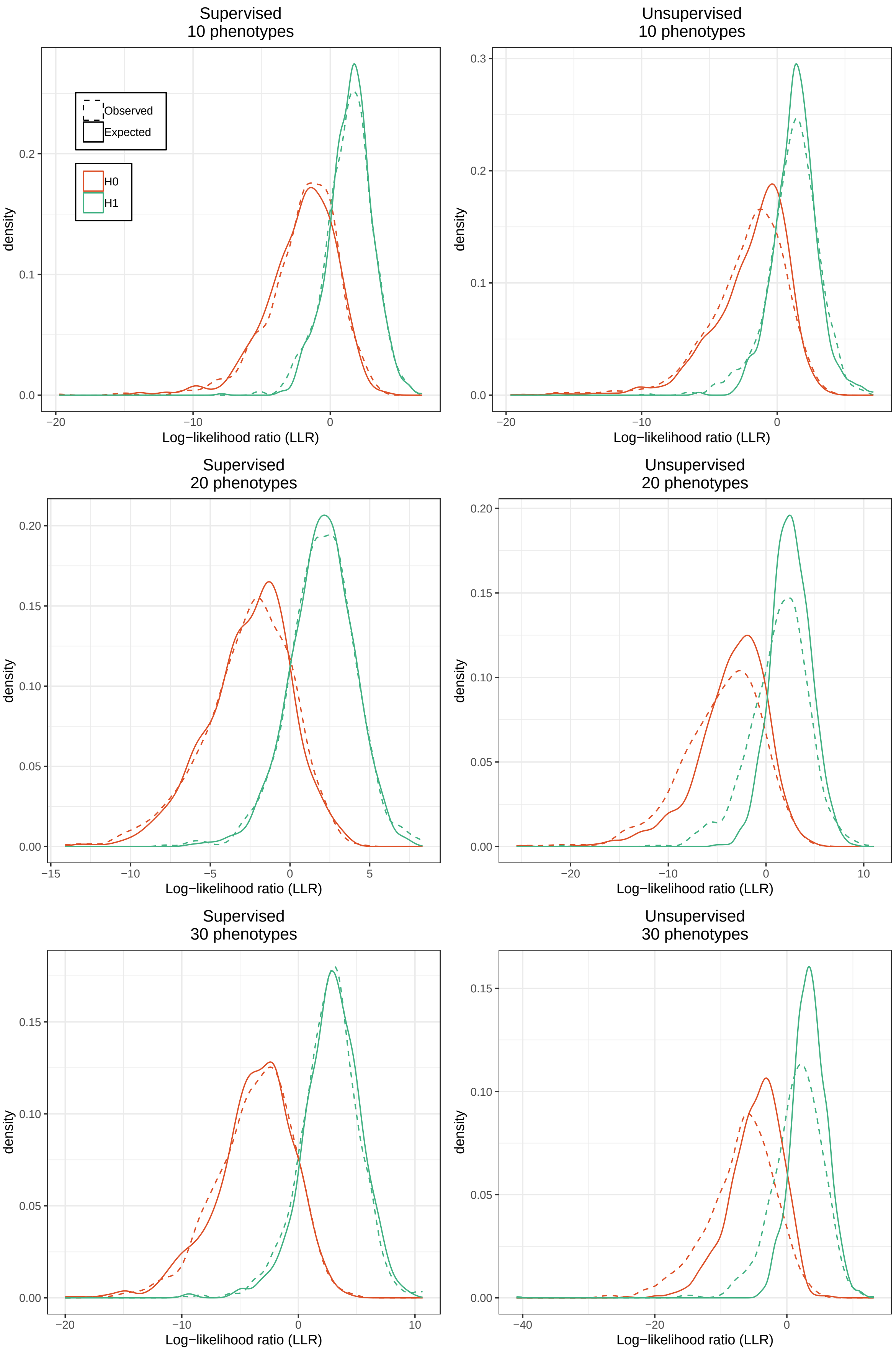
Supervised
10 phenotypes
Unsupervised
10 phenotypes
Observed
Expected
H0
H1
density
Log−likelihood ratio (LLR)
Supervised
20 phenotypes
Unsupervised
20 phenotypes
Supervised
30 phenotypes
Unsupervised
30 phenotypes

**Figure S2. AUPRC for match and mismatch inference.** Comparison of the area under the precision–recall curve (AUPRC) for match and mismatch inference using different number of phenotypes (from 30 to 40 phenotypes). **A** Results with the supervised approach. **B** Results with the unsupervised approach.

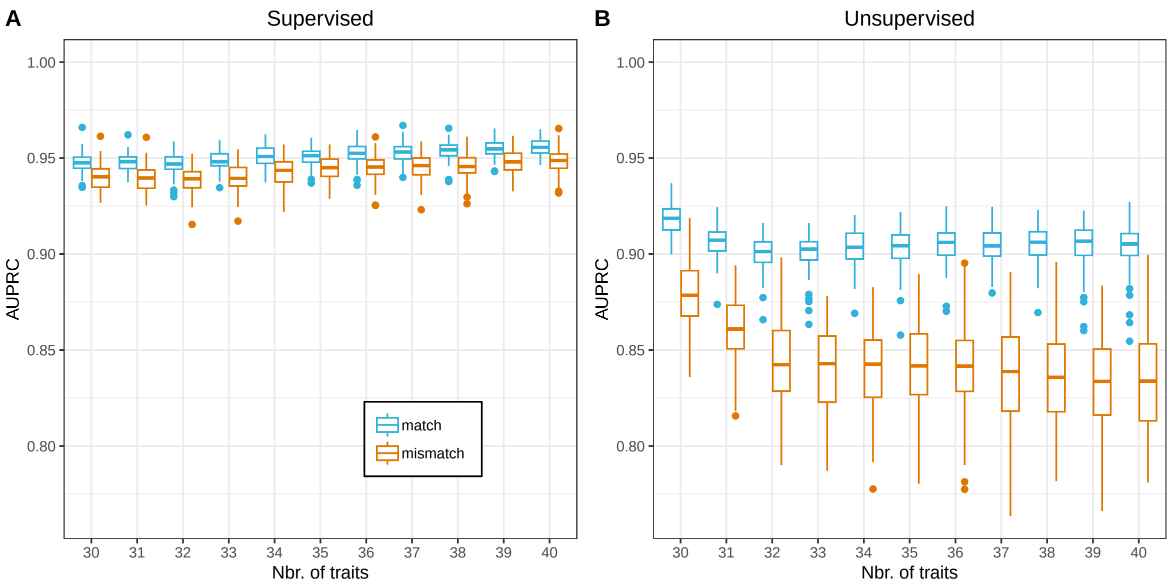


**Figure S3. AUPRC for match and mismatch inference excluding BMI and Immature reticulocyte fraction.** Comparison of the area under the precision–recall curve (AUPRC) for match and mismatch inference using different number of phenotypes (from 30 to 38 phenotypes) when excluding BMI and Immature reticuolcyte fraction. **A** Results with the supervised approach. **B** Results with the unsupervised approach.

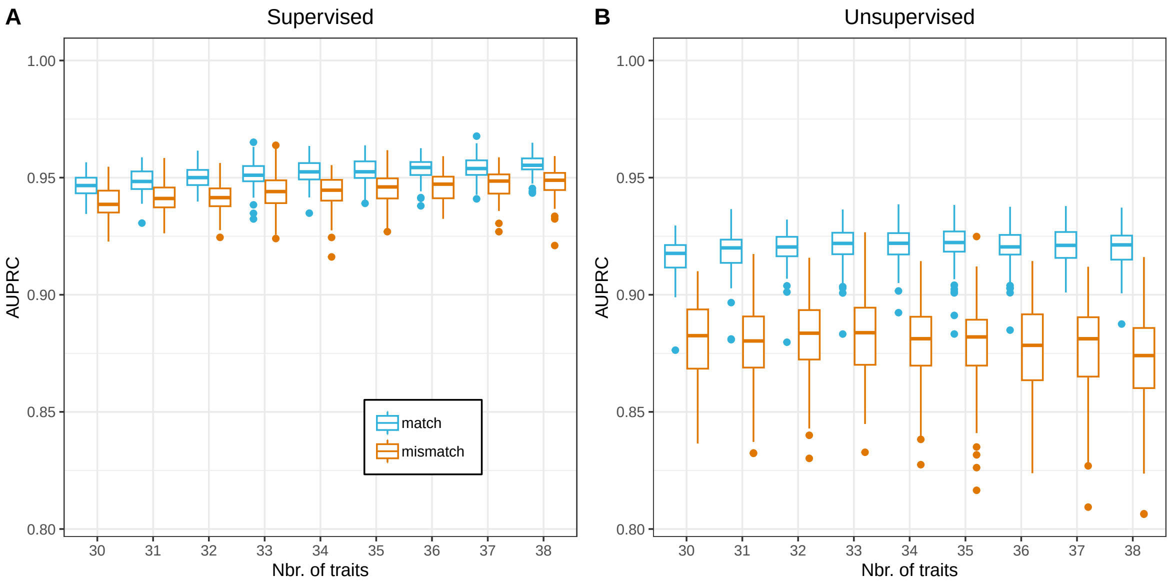


**Figure S4. Effect of using the genetic correlation as the environmental correlation in the unsupervised method**. Average precision recall curve for the supervised (light blue), unsupervised (dark blue) and unsupervised approach incorporating the environmental correlation estimated by the supervised approach (red). A, B, C and D correspond to results using different number of traits (see plot title).

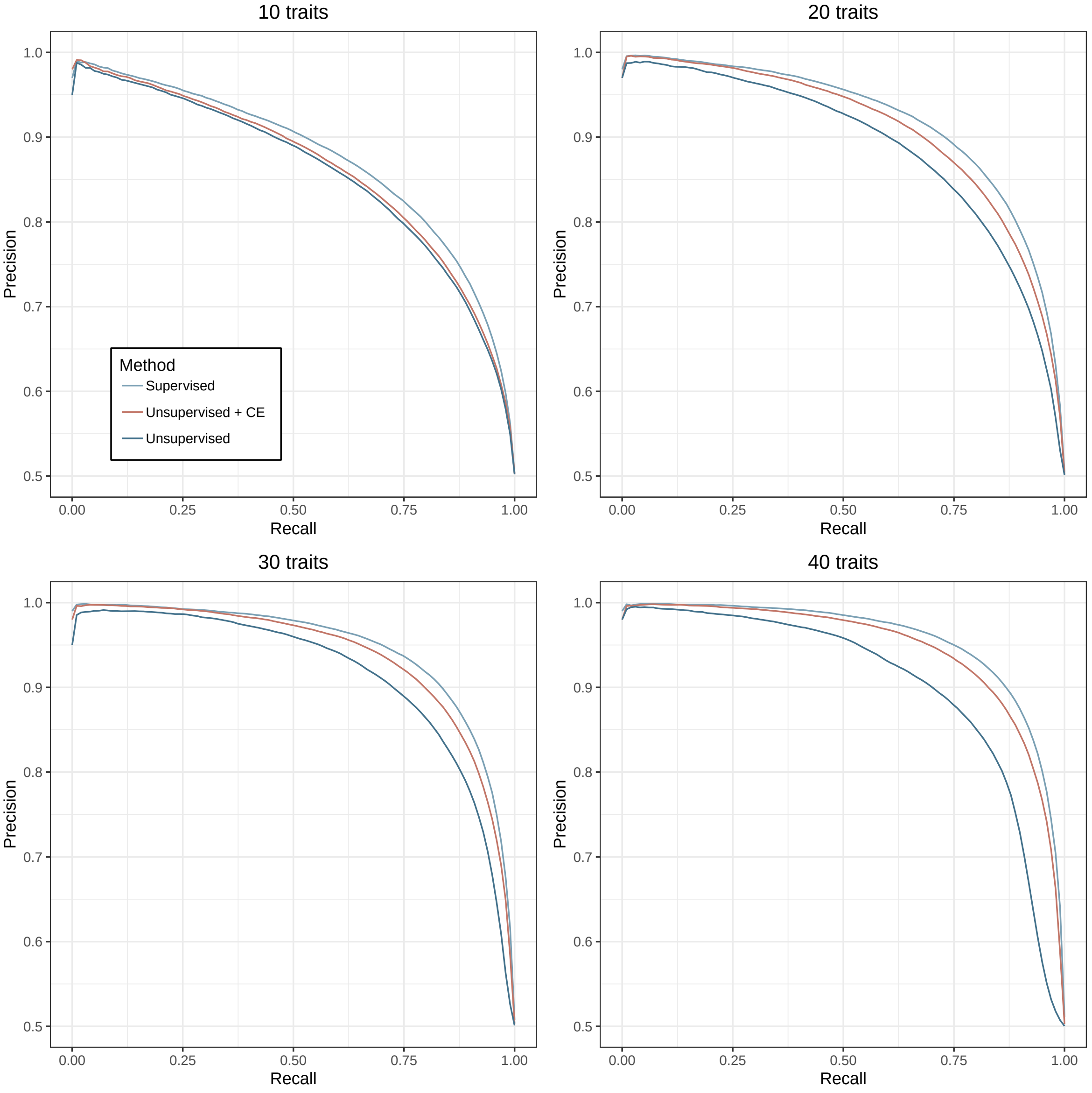


**Figure S5. Computational efficiency**. Comparison of computational time (**A,C**) and memory usage (**B,D**) for IDEFIX (blue) and our method (red). Panels (**A,B**) show results as a function of the number of phenotypes, while panels (**C,D**) show results as a function of the number of individuals in the test set. Bar heights represent the mean time/memory across 10 replicates. Error bars represent the minimum and maximum values across replicates.

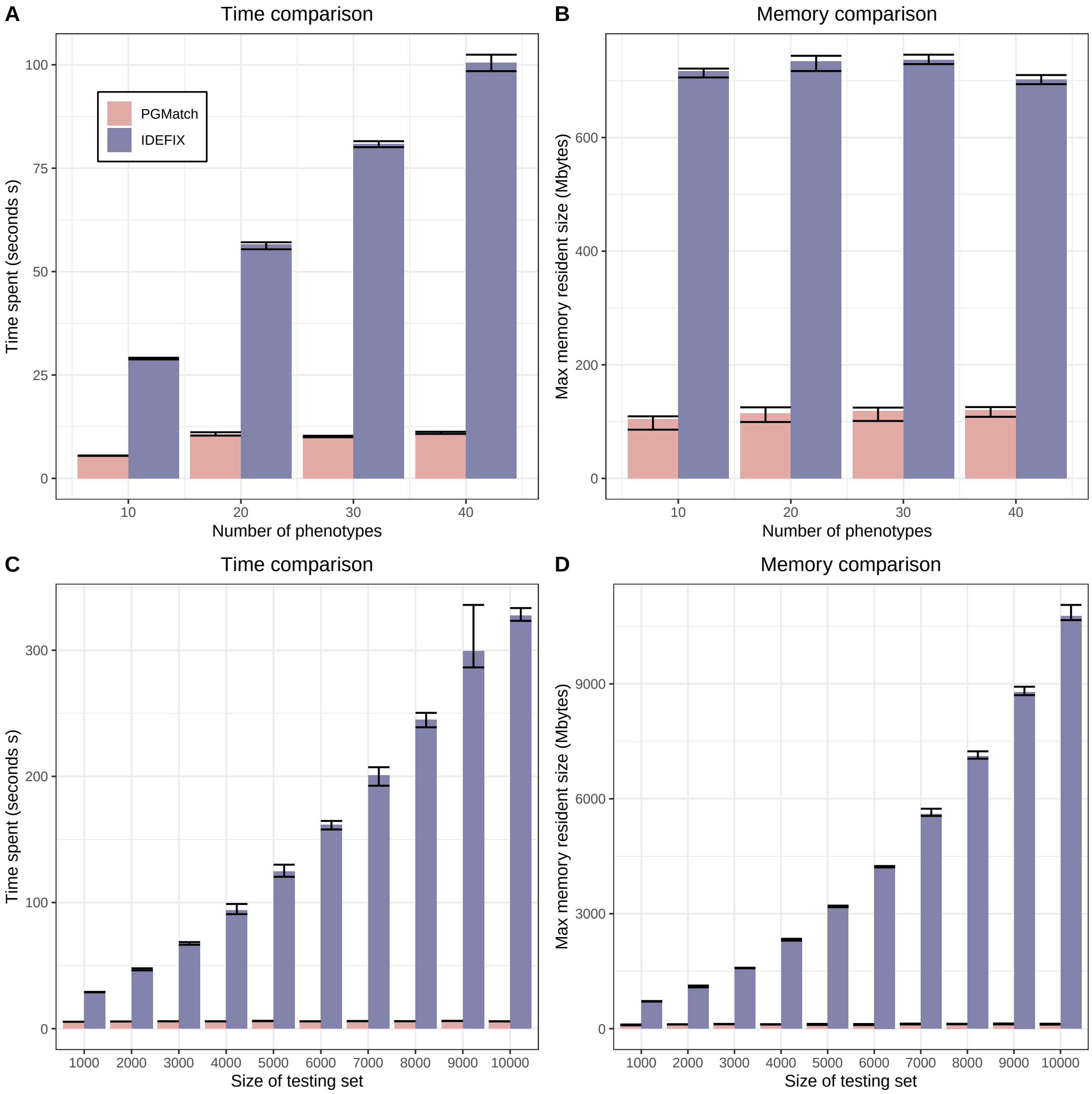


**Figure S6. Impact of phenotype-specific genetic variance.** Precision–recall curves for PGMatch and IDEFIX under different distributions of the proportion of variance explained by PGS ($r^2$). Colors indicate the $r^2$ distribution across phenotypes: Scenario 1 ($r^2 = 0.5$ for all phenotypes), Scenario 2 ($r^2 = 0.05$ for all phenotypes), Scenario 3 ($r^2 = 0.5$ for half of the phenotypes and $r^2 = 0.05$ for the other half), and Scenario 4 ($r^2 = 0.5$ for a single phenotype and $r^2 = 0.05$ for the remaining phenotypes). Line types indicate the method: supervised PGMatch (dashed) and IDEFIX (solid). Results are averaged over 100 simulations.

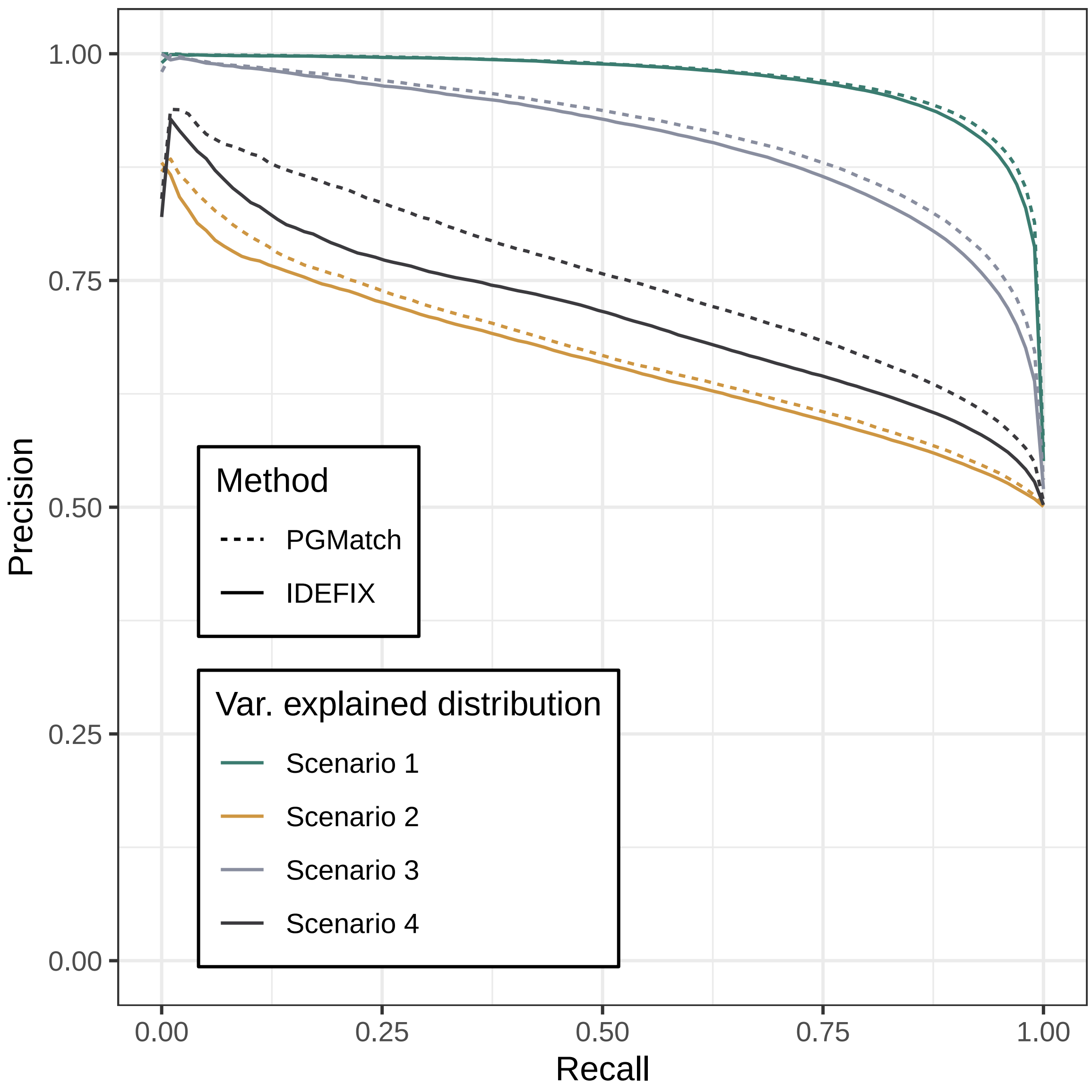


**Figure S7. Impact of PGS and environmental correlation.** Precision–recall curves for PGMatch and IDEFIX under different environmental ($\Sigma_e$) and PGS ($\Sigma_s$) correlations. Each plot correspond to a different distribution of the correlations: Scenario 1 ($\Sigma_e = 0$, $\Sigma_s = 0$); Scenario 2 ($\Sigma_e = 0$, $\Sigma_s = 0.5$); Scenario 3 ($\Sigma_e = 0.5$, $\Sigma_s = 0$), Scenario 4 ($\Sigma_e = 0.5$, $\Sigma_s = 0.5$) and Scenario 5 (heterogeneous correlations where half of the phenotypes have $\Sigma_e = \Sigma_s = 0.5$ and half have $\Sigma_e = \Sigma_s = 0$). Line types indicate the method: PGMatch (dashed) and IDEFIX (solid). Results are averaged over 100 simulations.

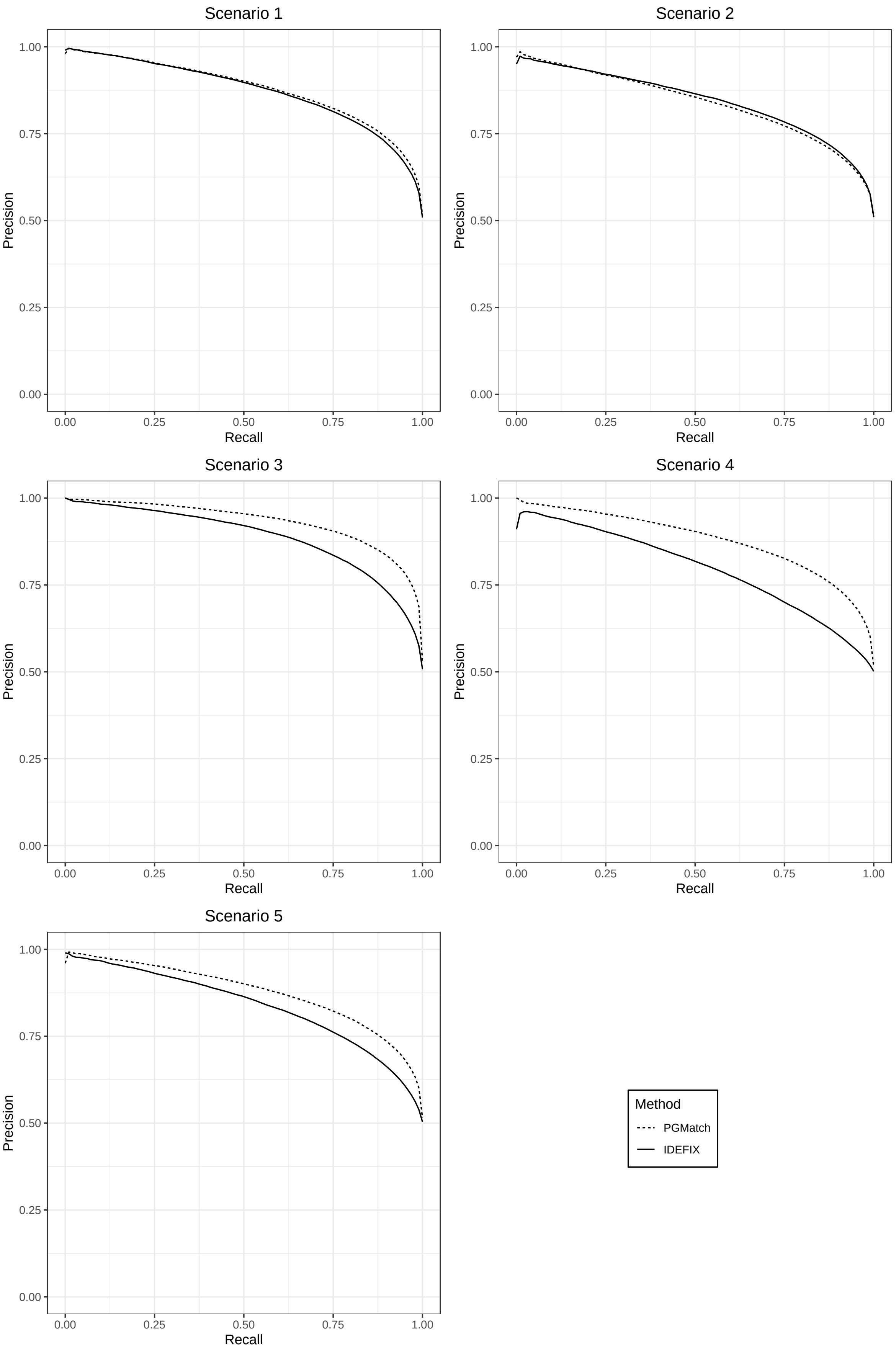
Scenario 1
Scenario 2
Scenario 3
Scenario 4
Scenario 5
Precision
Recall
1.00
0.75
0.50
0.25
0.00
Method
PGMatch
IDEFIX

**Figure S8. Inference of ABO status using re-identification by phenotypic prediction. A, B.** Pr(OO) in OO carriers and non-carriers using the supervised and unsupervised approach. The results of one-sided t-tests are shown above the connecting bars. **C, D.** Precision and recall for the inference of OO carriers using different Pr(OO) thresholds. The thick blue line represents the mean precision at a given recall level across 100 experiments. Ribbons represent the 95% bootstrap percentile interval of precision across experiments.

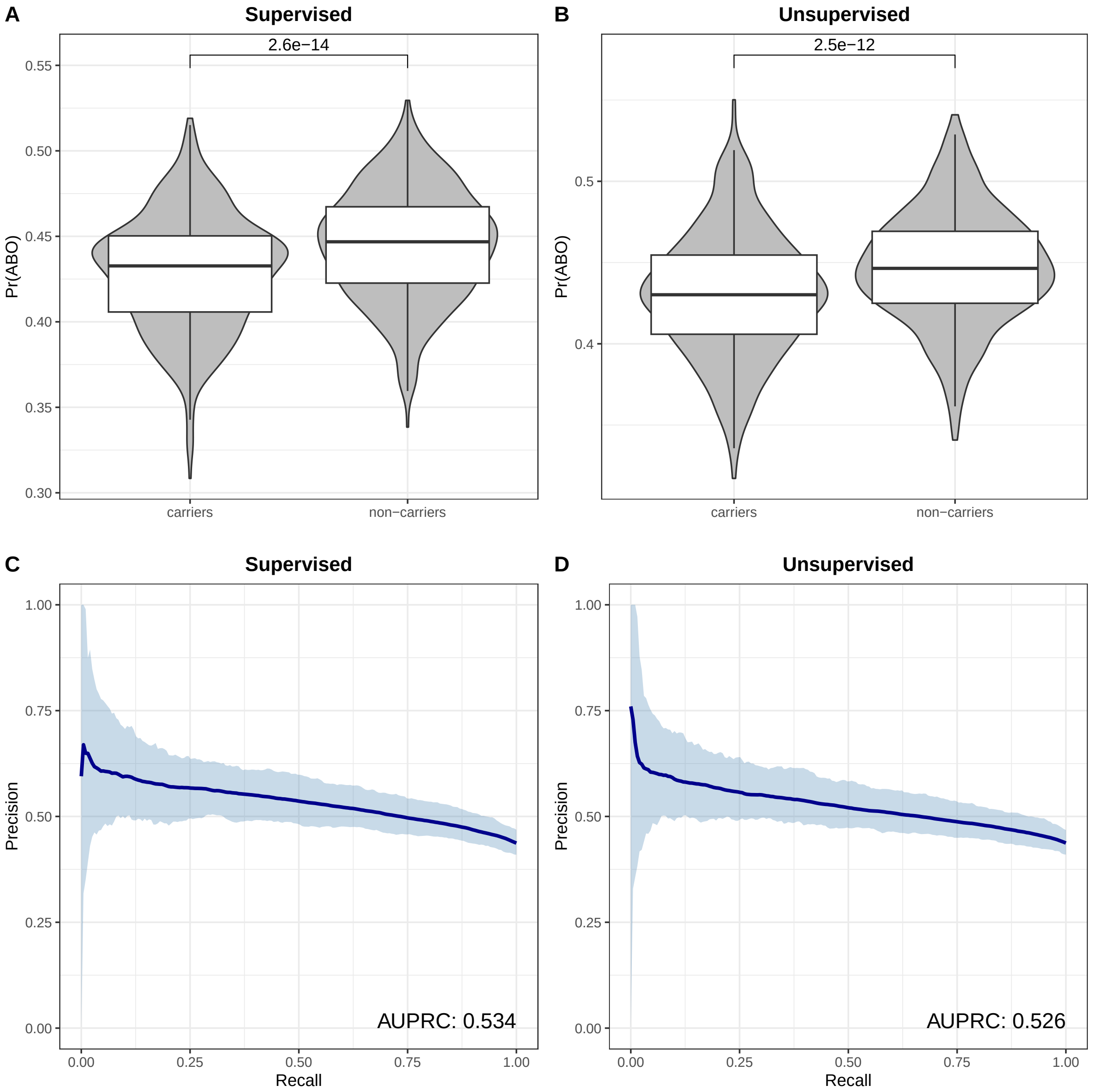


**Figure S9. Inference of FUT2 status using re-identification by phenotypic prediction. A, B.** Pr(FUT2) in FUT2 carriers and non-carriers using the supervised and unsupervised approach. The results of one-sided t-tests are shown above the connecting bars. **C, D.** Precision and recall for the inference of FUT2 carriers using different Pr(FUT2) thresholds. The thick blue line represents the mean precision at a given recall level across 100 experiments. Ribbons represent the 95% bootstrap percentile interval of precision across experiments.

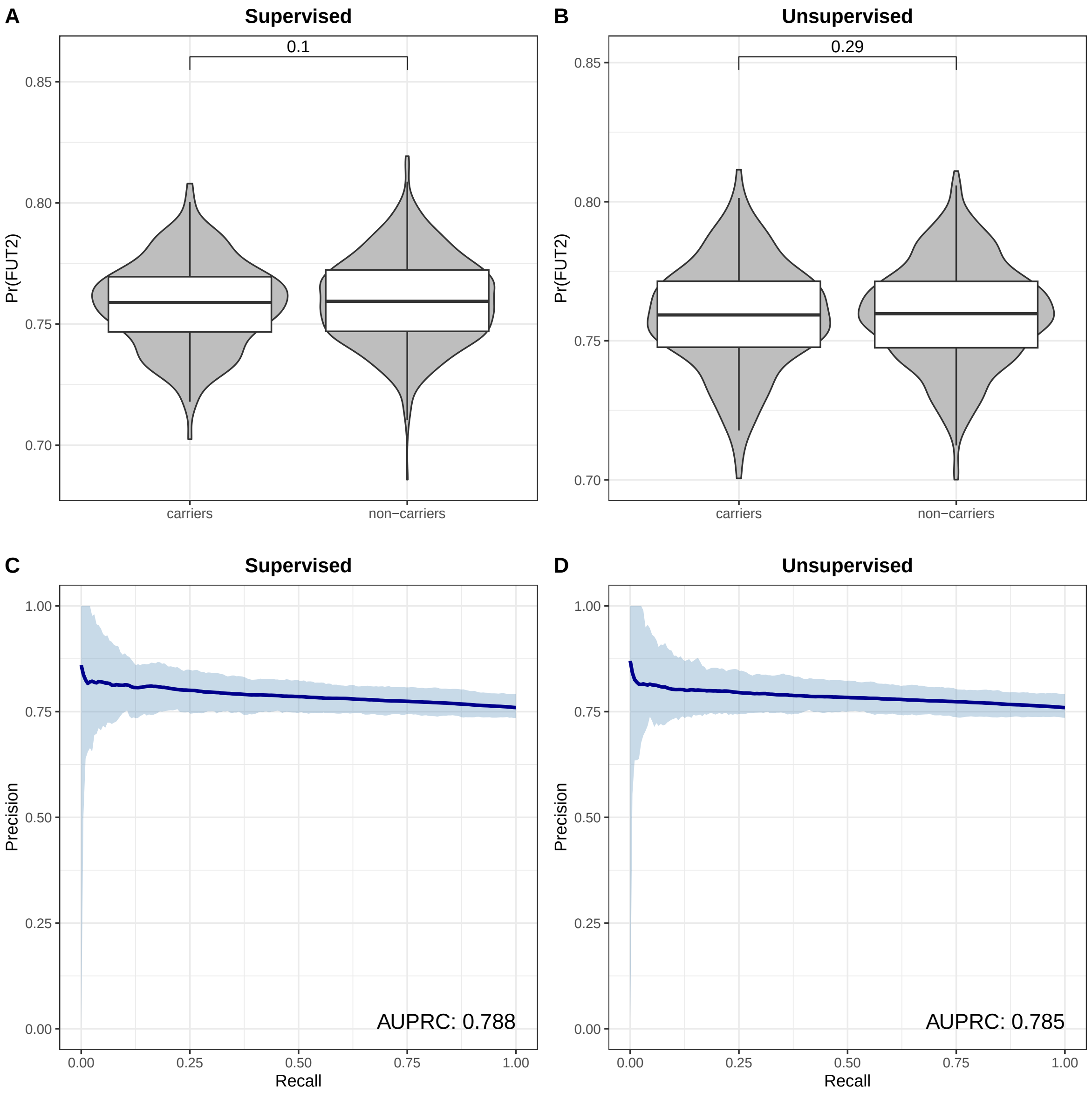


**Figure S10. Inference of LCT status using re-identification by phenotypic prediction. A, B.** Pr(LCT) in LCT carriers and non-carriers using the supervised and unsupervised approach. The results of one-sided t-tests are shown above the connecting bars. **C, D.** Precision and recall for the inference of LCT carriers using different Pr(LCT) thresholds. The thick blue line represents the mean precision at a given recall level across 100 experiments. Ribbons represent the 95% bootstrap percentile interval of precision across experiments.

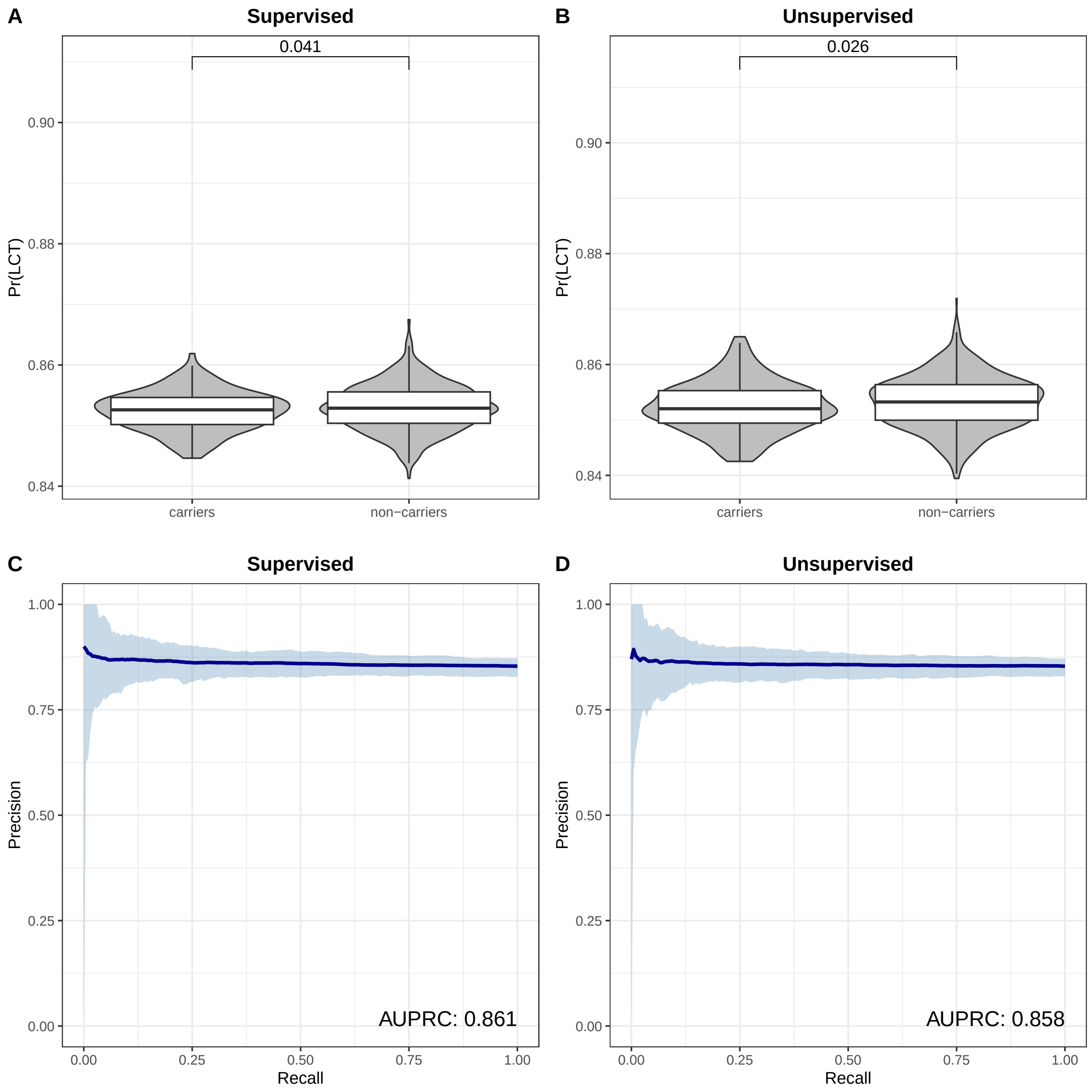


**Figure S11. Inference of *HLA-DQB1*06:02* status using re-identification by phenotypic prediction. A, B.** Pr($DBQ1\_602$) in *HLA-DQB1*06:02* carriers and non-carriers using the supervised and unsupervised approach. The results of one-sided t-tests are shown above the connecting bars. **C, D.** Precision and recall for the inference of *HLA-DQB1*06:02* carriers using different Pr($DBQ1\_602$) thresholds. The thick blue line represents the mean precision at a given recall level across 100 experiments. Ribbons represent the 95% bootstrap percentile interval of precision across experiments.

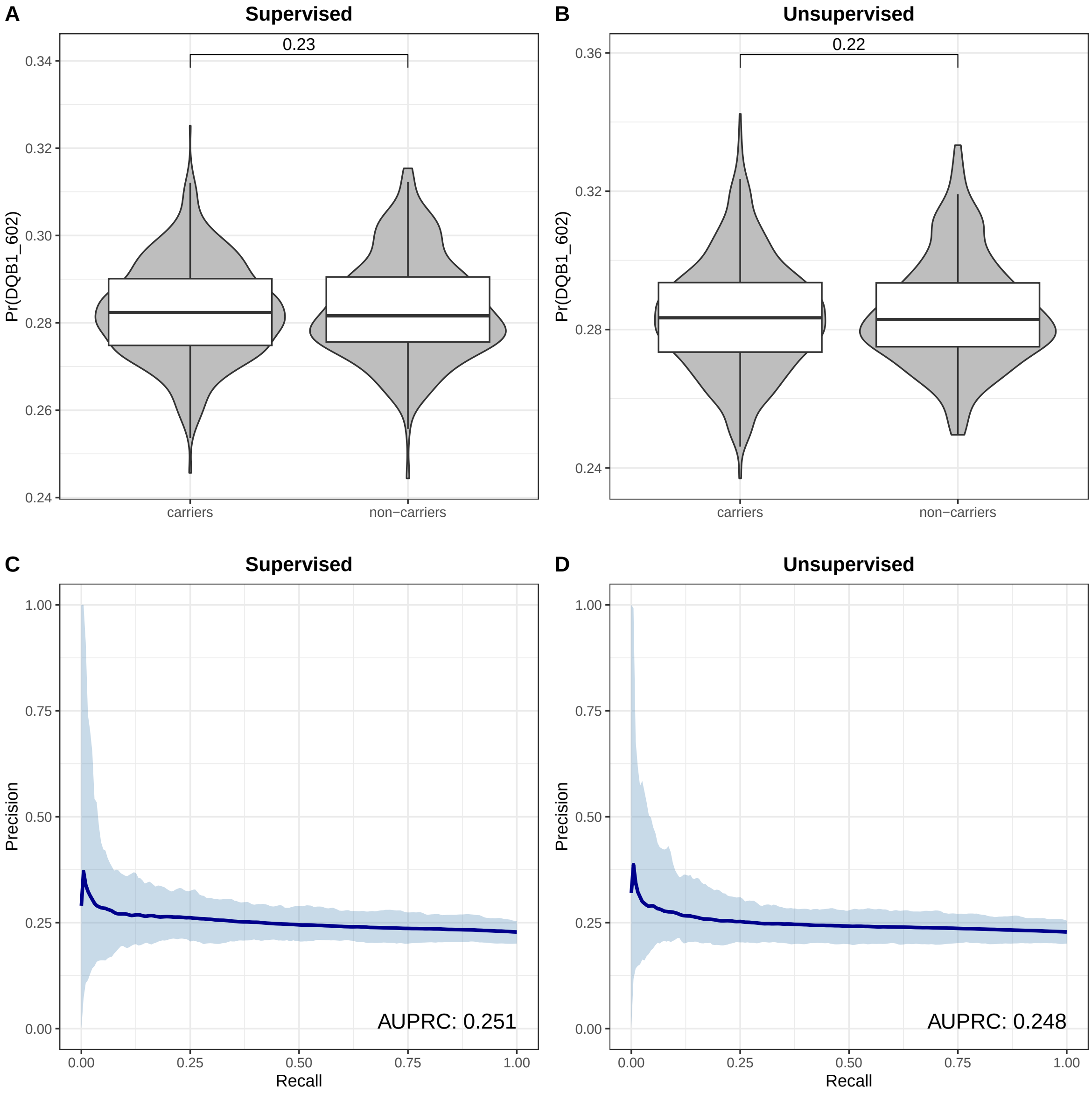


**Figure S12. Inference of genetic variant status using generalized linear models.** Precision–recall curves for the classification of genetic variant carriers using GLMs. Each panel corresponds to a different genetic variant: (A) APOE, (B) ABO, (C) FUT2, (D) LCT, and (E) *HLA-DQB1*06:02* Thick lines represent the mean precision across 100 bootstrap replicates, and ribbons indicate the 95% bootstrap percentile interval of precision across experiments.

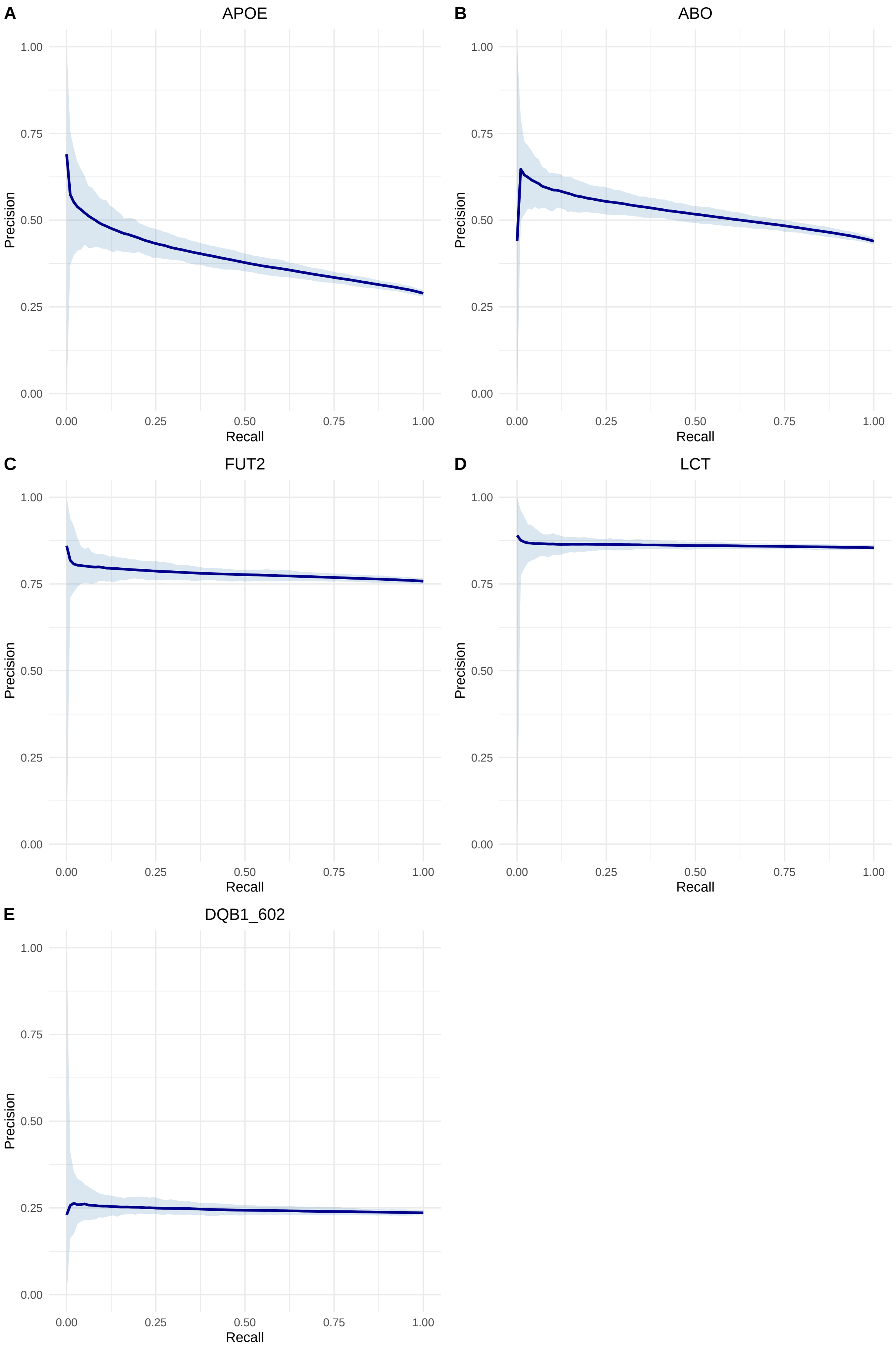
A
APOE
B
ABO
C
FUT2
D
LCT
E
DQB1_602
Precision
Recall
0.00
0.25
0.50
0.75
1.00

**Figure S13. Inferring if an individual is part of a biobank.** Mean (**A**) and Maximum (**B**) of matching probabilities between 1 000 individuals and 100 000 genomes in a biobank, depending on whether they were part of the biobank or not.

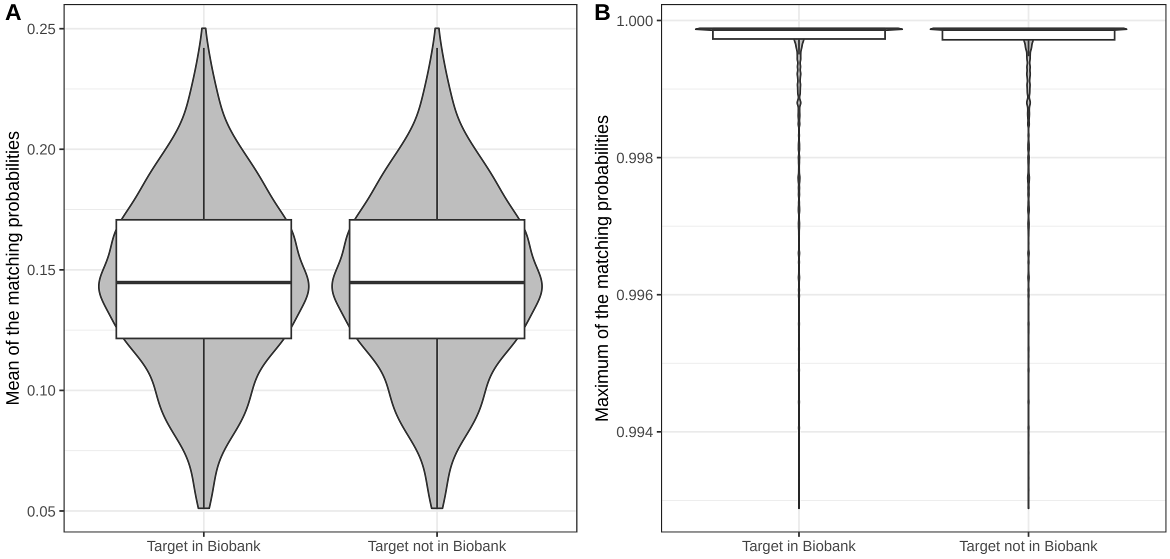
